# Undergraduates’ reactions to errors mediates the association between growth mindset and study strategies

**DOI:** 10.1101/2023.09.25.559345

**Authors:** Anastasia Chouvalova, Anisha S. Navlekar, Devin J. Mills, Mikayla Adams, Sami Daye, Fatima De Anda, Lisa B. Limeri

## Abstract

**Background:** Students employ a variety of study strategies to learn and master content in their courses. Strategies vary widely in their effectiveness for promoting deep, long-term learning, yet most students use ineffective strategies frequently. Efforts to educate students about effective study strategies have revealed that knowledge about effective strategies is by itself insufficient for encouraging widespread and lasting changes. An important next step is to uncover factors that influence the decisions students make about study strategy use. We explored the association between beliefs about intelligence (mindset, universality, and brilliance) and study strategies. The most effective study strategies are error-prone, and beliefs about intelligence carry implications for whether errors are a normal and even beneficial part of the learning process (e.g., growth mindset) or signs of insufficient intelligence (e.g., fixed mindset). Therefore, we hypothesized that beliefs about and reactions to errors would mediate a relationship between beliefs about intelligence and study strategies. We tested this hypothesis by surveying 345 undergraduates enrolled in an introductory biology class at a public, research-active university in northwestern United States.

**Results:** Confirmatory factor analysis indicated that the internal structure of all measures functioned as expected in our sample. We fit a structural equation model to evaluate our hypothesized model. We found that mindset, but not universality nor brilliance, predicts variance in both beliefs about errors and reactions to errors. In turn, adaptive reactions to errors (but not beliefs about errors) are associated with the use of highly effective study strategies and spacing study sessions. There was a significant indirect relationship between growth mindset and spacing of study sessions.

**Conclusions:** Our results provide evidence for a mechanism explaining the association between students’ mindset beliefs and academic outcomes: believing that intelligence is improvable is associated with more adaptive reactions to making errors, which correlates with choosing more error-prone and therefore more effective study strategies. Future interventions aimed at improving students’ study strategies may be more effective if they simultaneously target reacting adaptively to errors and emphasize that intelligence is improvable.

## INTRODUCTION

Undergraduates must engage with their class content outside the classroom (i.e., study) in order to master the material. In undergraduate contexts, the manner and timing with which students engage with the material outside of class is unstructured and managed by the student. These decisions are highly consequential for students’ mastery of material and their academic outcomes. Yet, students rarely receive explicit training or guidance about the most effective ways to study and there is great variation in student study strategies and patterns (Hartwig & Dunlosky, 2012; Karpicke et al., 2009; McDaniel & Einstein, 2020). Research in cognitive psychology and neuroscience has generated an extensive body of knowledge about the neurobiological mechanisms underpinning learning and the consequent effectiveness of different study strategies (Ambrose et al., 2010; Brown et al., 2014; Dunlosky et al., 2013; McGuire, 2018). When students use ineffective study strategies, they undermine their academic potential (Plant et al., 2015). Study strategies influence many features of student learning, including the amount of time students spend studying, how effectively they use this time, depth of conceptual understanding, and content mastery achieved (e.g., Pressley et al., 1987; Runquist, 1983). Ineffective study strategies cost time and do not effectively help students learn. Plant and colleagues (2005) surveyed undergraduates and asked them to log their study time and activities. They found that the amount of time students spent studying only influenced GPA when the effectiveness of the study strategies was accounted for (Plant et al., 2005). Thus, academic outcomes could potentially be improved by helping students learn about and ultimately adopt efficient study strategies. This may be particularly critical for students who face time constraints in working towards their educational goals, such as those with responsibilities to care for family and those who work outside of their studies.

### Study strategy effectiveness

There are many learning activities students engage in when they study, and these different study strategies vary substantially in how effectively they promote deep, long-term learning (Dunlosky et al., 2013). Broadly speaking, highly effective strategies are those that involve retrieving information from long-term memory, referred to as retrieval or recall practice (Carpenter et al., 2008; Dunlosky et al., 2013; Roediger & Karpicke 2006). These strategies are also referred to as deep learning strategies (e.g., Floyd et al., 2009; Sabah et al., 2023) because they are associated with forming connections between new and previous knowledge, reflecting on knowledge, and creating understanding (Biggs, 1987; Biggs & Tang, 2011; Marton and Säljö, 1976). In contrast, strategies that involve more passive engagement with material, such as re-reading and highlighting, are less effective at promoting learning (Carpenter et al., 2008; Dunlosky et al., 2013; Roediger & Karpicke 2006). These strategies are sometimes called surface learning strategies (e.g., Floyd et al., 2009; Good et al., 2013) because they contribute to a shallow learning approach – that is, fixating on trivial details and memorizing them, and lacking profound reflection (Schmeck, 1988).

The timing of study sessions, or study pattern, also impacts the effectiveness of studying (Dunlosky et al., 2013). Massing (also called cramming), where students attempt to do a large amount of studying in a short time period (especially the night before a test), is a common practice. However, spacing out study sessions (also called distributed practice) is more effective for learning and retention (Cepeda et al., 2006; Kornell, 2009). When students engage in massed practice, they are working primarily from their short-term, or working, memory. Spacing study strategies is more effective because the passage of time allows the information to leave short-term memory and forces recall to activate and strengthen neural pathways in long-term memory (Ambrose et al., 2010).

The study pattern (i.e., timing) and study strategies that students use relate to their academic outcomes. Rodriguez and colleagues (2018) surveyed over 1,200 undergraduates in a molecular biology course and found that those who reported self-testing and spacing had higher course grades. Hartwig and Dunlosky (2012) surveyed 324 undergraduates about their study strategies and academic outcomes. They found that students who reported self-testing also reported higher GPA and that students who reported massing (vs. spacing) their study sessions also reported using fewer overall study strategies. Williams and colleagues (2021) surveyed incoming undergraduates about their study strategies and found that those who reported self-testing also had significantly higher admissions GPA. Ewell and colleagues (2023) surveyed biology undergraduates in three different courses and found that using effective study strategies was associated with higher exam performance.

### Undergraduates’ study strategies

Literature characterizing undergraduates’ study behaviors indicates that students tend to pack their studying time with inefficient study strategies while dedicating a disproportionately low amount of time to more effective strategies (Carrier, 2003; Hartwig & Dunlosky, 2012; Karpicke et al., 2009; Morehead et al., 2016; Rea et al., 2022; Van Etten et al., 1997). A meta-analysis of students’ study strategies indicates that students most commonly report studying by re-reading and highlighting, which are less effective strategies (Miyatsu et al., 2018). Hartwig and Dunlosky (2012) surveyed 324 undergraduates (78% freshmen) and found that 66% of respondents reread their materials and 66% tend to cram the prior night. Morehead et al. (2016) found similar patterns in a sample of 300 undergraduates: 67% of participants reported rereading their materials and 53% of them tended to cram. In Karpicke and colleagues’ (2009) survey of 177 undergraduates, re-reading was the most commonly reported study strategy. Studies of community college biology students report similar patterns; students (n=52) reported re-reading material most frequently, followed by using flashcards and then underlining or highlighting while reading (Vemu et al., 2022). When students engage in inefficient study strategies, they are potentially suffering opportunity costs by spending time learning inefficiently.

### Errors are a critical part of effective learning

Generating errors is central to the process of learning (Mera et al., 2022). More specifically, generative learning refers to the process of forming deep connections between the new content and existing knowledge schemas, as well as rearrangement of neural networks (Fiorella, 2023; Fiorella & Mayer, 2016). Making sense of new information involves the generation of errors and of learning from these errors to address misconceptions, facilitate conceptual change, and reinforce correct conceptions. Unsurprisingly, many of the effective learning strategies linked to generative learning are the same, including self-testing, self-explanation, and creating visualizations, all of which are essentially sense-making strategies that result in deep learning (Fiorella & Mayer, 2016). Compared to passively reading materials, errorful learning paired with corrective feedback is more beneficial to student learning and retention (Mera et al., 2022; Overman et al., 2021).

Cognitive psychology research indicates that strategies that feel easy to students, such as re-reading and highlighting, feel effective to students, yet are ineffective for actual learning (Deslauriers et al., 2019; Dunlosky et al., 2013; Kirk-Johnson et al., 2019; Macaluso et al., 2022). This false feeling of effectiveness is referred to as the fluency effect (Deslauriers et al., 2019; Macaluso et al., 2022). In contrast, strategies that are highly effective, such as self-testing and spacing, involve students making errors and can feel frustrating (Deslauriers et al., 2019; Dunlosky et al., 2013; Kirk-Johnson et al., 2019; Macaluso et al., 2022). Because they feel frustrating and are error-prone, these effective strategies can induce anxieties, negatively impact self-efficacy, and be perceived as costly (Rea et al., 2022). The fluency effect explains that if students passively engage with their material many times, which is how many students choose to study (Carrier, 2003; Hartwig & Dunlosky, 2012; Karpicke et al., 2009; Miyatsu et al., 2018; Morehead et al., 2016; Rea et al., 2022; Van Etten et al., 1997), they develop an *illusion of competency* (Bjork, 1999; Kelley & Jacoby, 1996; Koriat & Bjork, 2005; Nyland & Sawarynski, 2017).

Numerous studies with undergraduates have supported the *misinterpreted effort hypothesis*—that students misinterpret expending greater effort as a sign of lower efficacy (Kirk-Johnson et al., 2019). Kirk-Johnson and colleagues (2019) conducted a series of four survey-based studies in which they demonstrated that: (1) students viewed effective study strategies as more effortful and worse for learning, (2) mental effort indirectly affected strategy choice via perceived learning, and (3) learners interpret high mental effort required by effective strategies as a sign of poor learning. Deslauriers and colleagues (2019) supported this hypothesis in a classroom-based experiment. They taught introductory physics content to undergraduates using active-learning and traditional didactic methods. They found that students taught by active learning learned more but perceived that they learned less than when they were taught by didactic methods. Similarly, Kornell (2009) found that students who studied word-pairs learned more effectively with spaced practice, yet 72% of the participants believed that cramming was more effective.

There is also evidence that students believe massing study sessions is more effective than spacing them out because massing makes the study activities feel easier (Baddeley & Longman, 1978). Because spacing forces recall to activate long-term memory, it causes errors and can reduce performance levels during the study sessions (Bjork, 1999). Therefore, students often believe that cramming is more effective because it can result in higher short-term performance, which they mistake for long-term learning (Bjork, 1999).

Educators have long acknowledged the productive role of making errors in learning and embedded them intentionally in instructional strategies. Bjork (1999) coined the term “desirable difficulties” to describe learning conditions that increase the cognitive effort required and thus lead to deeper and more long-lasting learning. Productive failure is a pedagogical approach that intentionally embeds desirable difficulties into learning activities by tasking students to solve difficult, complex problems before they have been taught all the necessary tools and information (Kapur, 2008; Kapur & Bielaczyc, 2012). Struggling with these initial challenging problems helps students better learn and remember the information when they are subsequently taught it and increases later performance on similar problems (Kapur, 2008).

### Study strategy interventions

Some educators and researchers have attempted to leverage findings from cognitive science to help students adopt more effective study strategies and thus reap better academic outcomes. Rodriguez and colleagues (2018) conducted light-touch interventions in an introductory biology course, consisting of a 10-min lecture about the benefits of self-testing and spacing and weekly reminders throughout the term that students should implement these strategies. Compared to students in control sections, students who received this light-touch intervention reported using self-testing and spacing more often (Rodriguez et al., 2018). Vemu and colleagues (2022) conducted a similar intervention in a community college context, in which students received a brief lecture about effective study strategies and completed an assignment that prompted them to reflect on their study strategies after each exam. Students reported increasing using spacing and generating drawings as study strategies (Vemu et al., 2022).

However, not all study strategies interventions have been successful in shifting students’ behaviors and improving academic outcomes (Hattie et al., 1996; McDaniel & Einstein, 2020; Wang et al., 2023). McDaniel and Einstein (2020) note that most study strategy programs and interventions fail to improve students’ self-regulation and metacognition and that students fail to apply good study strategies across various content and contexts. They detail numerous barriers that students face in adopting effective strategies, including educational contexts focused on rote memorization that encourage surface-learning strategies, the lack of timely and accurate feedback available to studiers about the effectiveness of strategies they employ, and the unintuitiveness of effective study strategies (McDaniel & Einstein, 2020). Wang, Muenks, and Yan (2023) found that perceived cost was negatively associated with students’ reported use of retrieval practice as a study technique. One study found that over 80% of undergraduate psychology students reported that they do not follow their instructor’s study strategy suggestions (Kornell & Bjork, 2007). Another study of undergraduates found that students report understanding the benefits of spacing, but still massed their study sessions frequently and engaged in inefficient study strategies, like re-reading (Susser & McCabe, 2013). Studies of undergraduates’ metacognitive knowledge and skills found that students sometimes choose to use study strategies that they know are ineffective because they feel comforting during a stressful time (Dye & Stanton, 2017). In fact, in their study, a student likened using familiar, ineffective study strategies to eating comfort food (Dye & Stanton, 2017). Knowledge of the effectiveness of study strategies is not always sufficient for students to change their habits, as many students make plans to study using effective strategies which they are unable to follow through on (Stanton et al., 2015).

## THE CURRENT STUDY

A major challenge to improving students’ study strategies via interventions is that knowledge of study strategies, by itself, does not spark widespread change in students’ study behaviors (Rea et al., 2022; Stanton et al., 2021). Effectively encouraging students to adopt effective study strategies will require an approach informed by the factors that influence students’ decisions about how to study. Our study seeks to explore motivational factors underpinning students’ study strategy behaviors.

Since errors play a central role in the learning process, we hypothesize that students’ beliefs about errors and reactions to errors may influence how they choose to study. Prior work has explored how students approach and respond to errors while learning. Tulis and colleagues (2017) conceptualized and explored the beliefs students hold about learning from errors, called error-related beliefs. Positive beliefs about errors reflect the attitude that one can benefit from making errors in the learning process whereas negative beliefs about errors reflect the attitude that errors are harmful and should be avoided.

Dresel and colleagues (2013) characterized two types of reactions that students have towards errors: action-oriented and affective-motivational. They conceptualized action adaptivity of error reactions as how individuals change their course of action in response to making an error. For example, students may put in more effort next time and set goals, or on the contrary, disengage and invest less effort. Affective-motivational adaptivity of error reactions describes how making errors affects students’ motivation and emotions, and is associated with students’ persistence, performance, and effort (Dresel et al., 2013; Grassinger & Dresel, 2017; Kreutzmann et al., 2014). For example, a student may be more motivated and have just as much fun in the class if they make errors, or they become less motivated and find the class more threatening in the future.

To our knowledge, prior work has not connected error-related beliefs or reactions to errors to students’ choice of study strategies. Given the central role of errors in effective study strategies, **we hypothesize that positive beliefs about errors and positive reactions to errors would be associated with using more error-prone and effective study strategies, like spacing out study sessions and using self-testing, writing out notes from memory, and creating visuals as study techniques (Figure 1).**

**Figure 1.**
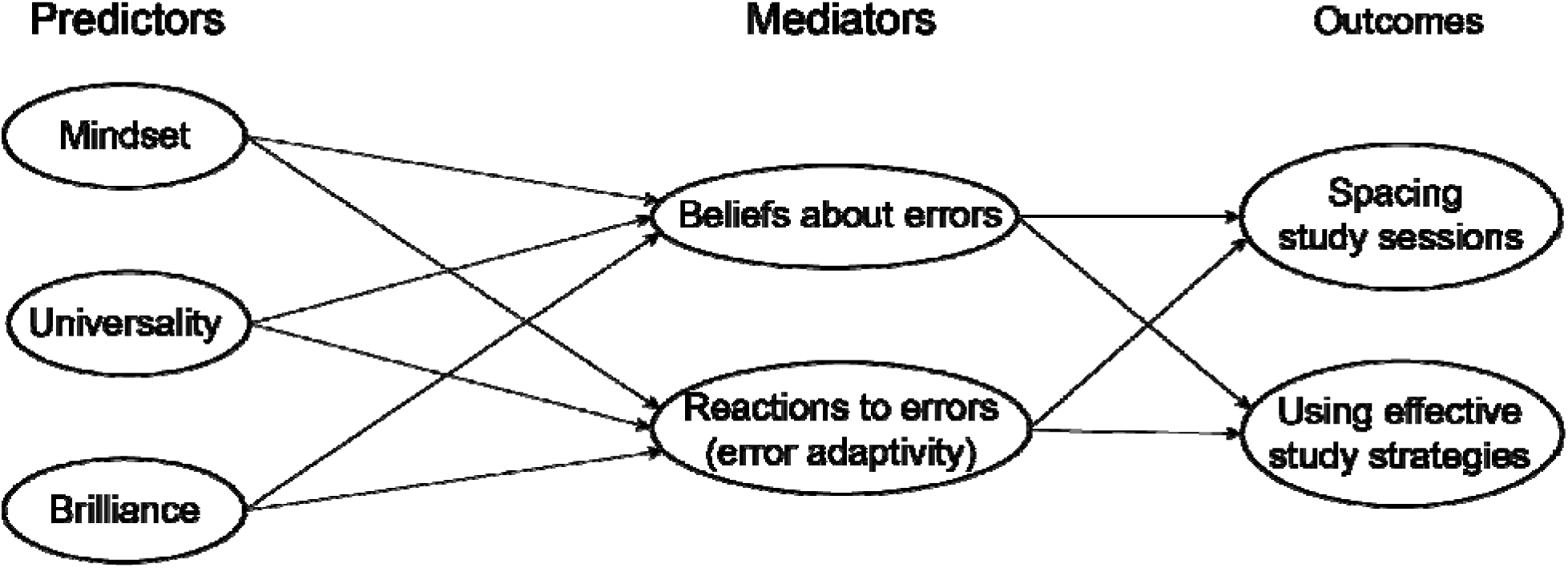
Hypothesized mediation model. We hypothesize that beliefs about abilities (mindset, universality, and brilliance) will influence attitudes towards errors and their responses to errors, which in turn will influence the study strategies students use.

Students’ attitudes towards and reactions to learning challenges have been tightly linked to students’ beliefs about the malleability of their intellectual abilities (termed mindset; Dweck, 1999). Mindset beliefs influence how students interpret their performance, including failures and negative feedback (Burnette et al., 2013; Haimovitz & Dweck, 2016; Hong et al., 1999; Schroder et al., 2014; Smiley et al., 2016; Sun & Huang, 2023; Tabibnia & Radecki, 2018; Waller & Papi, 2017; Yao & Zhu, 2022). In fact, Yeager and Dweck (2020) wrote in a recent commentary that “mindset is a theory about response to challenges and setbacks, not about academic achievement.” When students believe that their intellectual abilities are fixed and unchangeable (i.e., a fixed mindset), they interpret negative feedback or errors as an indictment of themselves and their ability, and as a personal threat (Burnette et al., 2013; Haimovitz & Dweck, 2016; Hong et al., 1999; Smiley et al., 2016). This leads students to withdraw from the threatening situation and reduces persistence (Burnette et al., 2013; Haimovitz & Dweck, 2016; Hong et al., 1999; Smiley et al., 2016). Thus, if students receive negative feedback about their mastery of the content while engaging in error-prone learning strategies, those with a fixed mindset might choose to discontinue using those strategies. In contrast, students who believe their abilities can improve (i.e., a growth mindset) see negative feedback about their performance as a normal part of the learning process and useful information to direct their efforts (Burnette et al., 2013; Haimovitz & Dweck, 2016; Hong et al., 1999; Smiley et al., 2016; Sun & Huang, 2023; Waller & Papi, 2017; Yao & Zhu, 2022). This trend is also supported by neuroimaging evidence.

Students with a growth mindset have shown higher error positivity waveform response, which is the neural correlate of being more aware of one’s own errors, more receptive of feedback, and having greater post-error accuracy (Mangels et al., 2006; Moser et al., 2011; Ng, 2018; Tirri & Kujala, 2016). In contrast, those with a fixed mindset perceive critical feedback as a threat to their self-perceptions of ability and fail to demonstrate deep processing of the feedback, leading to more subsequent errors (Mangels et al., 2006). These findings inform the so-called “social cognitive neuroscience” model (Mangels et al., 2006).

Based on these theoretical linkages and prior work, **we also hypothesized that students’ mindset beliefs would be associated with their beliefs about errors and reactions to errors (Figure 1).** Specifically, we hypothesize that students with growth mindset would hold positive beliefs about errors and react positively to errors, which should be related to their adoption of highly effective, error-prone study strategies. In contrast, students with a fixed mindset would hold more negative beliefs about errors and react negatively to making errors, and thus avoid using error-prone study strategies which are highly effective.

Mindset describes only one type of belief students hold about abilities (called “lay theories”). Two other lay theories have been described in educational literature more recently and are consequently less thoroughly studied. Universality refers to beliefs about whether everyone (universal belief) or only some individuals (non-universal belief) hold the potential to reach the highest levels of intellectual ability (Rattan et al., 2012; 2018). Brilliance belief refers to the belief about the degree to which an innate, high ability (brilliance) is required for success in a given field (Leslie et al., 2015; Meyer et al., 2015). Mindset, universality, and brilliance all describe beliefs regarding intellectual abilities and success. Prior work has shown that they are distinct beliefs and each uniquely predicts undergraduates’ psychosocial and academic outcomes (Limeri et al., 2023). This prior work revealed that universality and brilliance predicted variance in most psychosocial and academic outcomes related to mindset beyond what mindset alone was able to predict. While less work has been done on universality and brilliance beliefs, the theoretical underpinnings suggest they may relate to how students interpret and respond to errors. Students with a non-universal belief might view errors as a sign that they are not one of the special people with unique potential. Thus, they may view errors as threatening (similar to those with a fixed mindset) and have lower acceptance and poorer response to errors than those with a universal belief. Similarly, we reasoned that students with a strong brilliance belief may view errors as a sign that they do not have the brilliance required for success. This view would make errors threatening and may motivate students to avoid and have a negative attitude towards errors and react poorly to them.

Based on this theoretical connection, **we also hypothesized that brilliance and universality beliefs may influence students’ beliefs about and reactions to errors, and thus study strategy choices (Figure 1).** Specifically, we hypothesize that students with a universal belief would hold positive beliefs about errors and react positively to errors, which should be related to their adoption of highly effective, error-prone study strategies. In contrast, students endorsing the non-universal and brilliance beliefs would hold more negative beliefs about errors and react negatively to making errors, and thus avoid using error-prone study strategies which are highly effective.

The current study aims to clarify the relationships among beliefs about abilities, error-related beliefs and reactions, and study strategies. This work advances the field in several important ways. First, to our knowledge, this is the first study to investigate the potential role of beliefs about errors and reactions to errors as mediating factors between mindset and study strategies. Second, we use a newly developed, context-specific measure of mindset for undergraduate STEM students which has extensive evidence of validity and has shown higher predictive capacity than previously available measures (the ULTrA survey; Limeri et al., 2023; unpublished data). Third, this work expands the limited literature on the role of universality and brilliance beliefs – these are beliefs about abilities that have recently shown unique predictive utility in explaining various undergraduate psychosocial and academic outcomes (Limeri et al, 2023).

## METHODS

We conducted a quantitative study to test our hypothesized model (Figure 1). We surveyed undergraduate students (n = 345). The survey included measures of students’ beliefs about abilities (lay theories: mindset, universality, and brilliance), beliefs about errors, adaptivity to errors, and asked students to report the study strategies and patterns they use. The study protocol was reviewed and approved by the Texas Tech University Institutional Review Board (IRB2022-150). Consent was obtained from participants on the first page of the survey.

### Participants

We surveyed students enrolled in an introductory biology course at a public, selective, very high research activity, primarily white university located in Northwestern United States. The course enrolls ∼500 students who are typically about half first-year students and half upper-level students (∼25% second-year students, ∼25% third- and fourth-year students). Students were awarded a small amount of extra credit for completing the survey (or alternative assignment).

We received 413 responses to the survey. We removed 13 incomplete responses. The survey included two attention-check items (e.g., “This is a control question. Please select ‘Strongly disagree’”), located at approximately 1/3 and 2/3 of the way through the survey. Fifty-five responses failed to select the directed response to one or both checks and were removed, resulting in a final sample size of n = 345. Our sample was mostly white (75%), contained disproportionately more women (68%) than other genders, and included more continuing-generation students (70%) than first-generation students (Table 1).

**Table 1.**
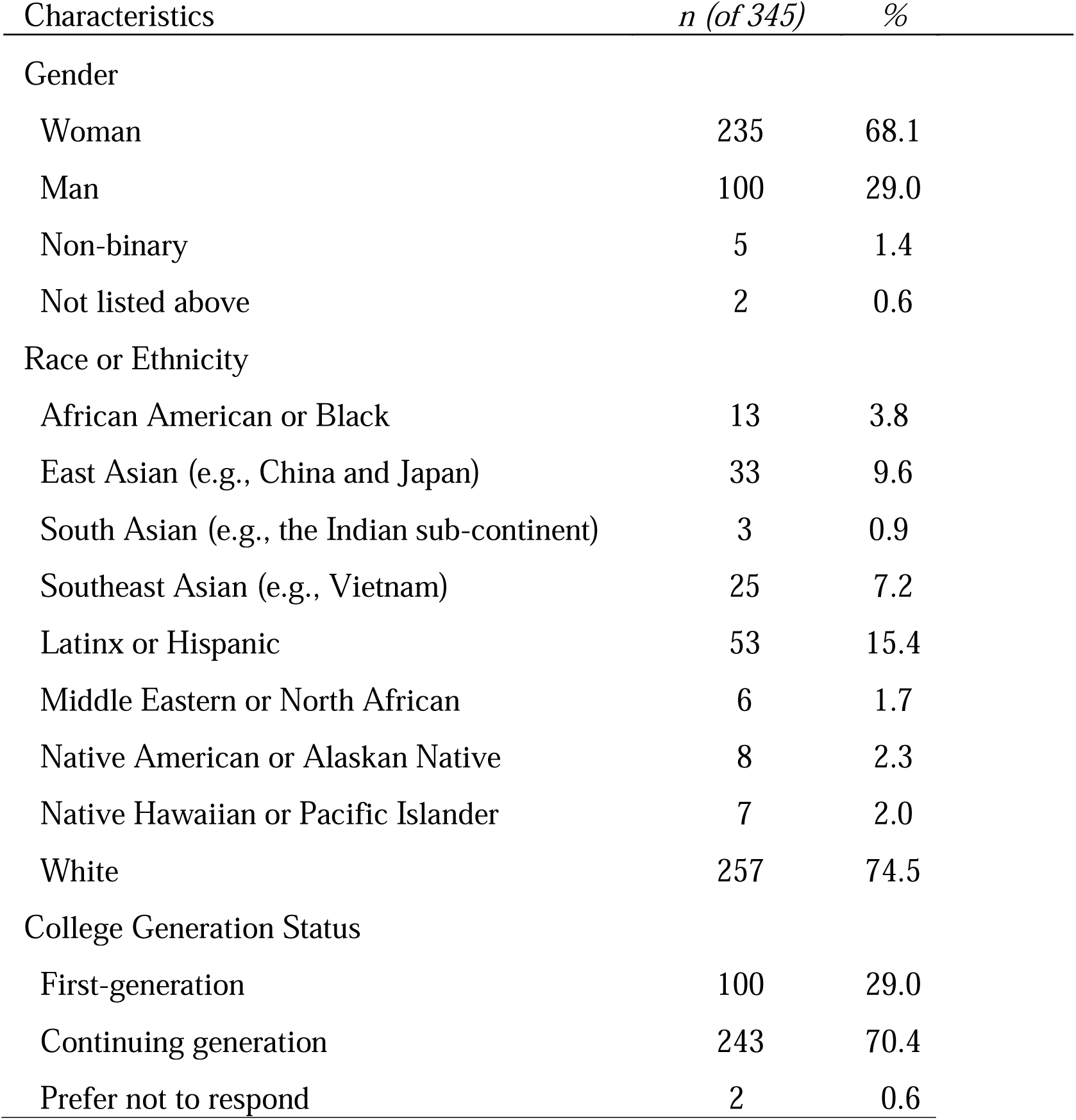
Demographics information of study participants.

| Characteristics | *n (of 345)* | *%* |
| --- | --- | --- |
| Gender |  |  |
| Woman | 235 | 68.1 |
| Man | 100 | 29.0 |
| Non-binary | 5 | 1.4 |
| Not listed above | 2 | 0.6 |
| Race or Ethnicity |  |  |
| African American or Black | 13 | 3.8 |
| East Asian (e.g., China and Japan) | 33 | 9.6 |
| South Asian (e.g., the Indian sub-continent) | 3 | 0.9 |
| Southeast Asian (e.g., Vietnam) | 25 | 7.2 |
| Latinx or Hispanic | 53 | 15.4 |
| Middle Eastern or North African | 6 | 1.7 |
| Native American or Alaskan Native | 8 | 2.3 |
| Native Hawaiian or Pacific Islander | 7 | 2.0 |
| White | 257 | 74.5 |
| College Generation Status |  |  |
| First-generation | 100 | 29.0 |
| Continuing generation | 243 | 70.4 |
| Prefer not to respond | 2 | 0.6 |

First-generation is defined as neither of the student’s parents/guardians having earned a four-year college degree. Continuing generation status is defined as having at least one parent/guardian who has earned a four-year college degree. Note that counts may not add up to the total number of participants if respondents declined to answer the question or selected more than one identity (e.g., for race/ethnicity).

### Data collection methods

The survey included measures of beliefs about abilities, beliefs about errors, error adaptivity, study strategies, and four demographic questions. The complete item set can be found in Supplemental Material Section 1.

#### Beliefs about abilities

We measured mindset, universality, and brilliance beliefs using the Undergraduate Lay Theories of Ability (ULTrA) Survey (Limeri et al., 2023). This survey was developed specifically for use with undergraduate STEM students and is supported by multiple types of validity evidence (Limeri et al., 2023). The ULTrA is a 25-item measure of growth mindset (5 items, e.g., “I can become excellent at applying knowledge to solve challenging problems”), fixed mindset (5 items, e.g., “At the end of college, my ability to analyze information will be at about the same level that it is now”), universal belief (5 items, e.g., “Anyone could become as effective at learning as highly successful STEM students”), non-universal belief (5 items, e.g., “Only people with a natural talent can become good enough at applying knowledge to solve the most difficult problems”), and brilliance belief (5 items, e.g., “Becoming a top student in STEM requires an innate talent that just can’t be taught”).

#### Beliefs about errors and error adaptivity

We used the measure that Tulis and colleagues (2017) developed and collected supporting validity evidence for use with undergraduates. This survey measures beliefs about learning from errors (5 items, e.g., “Errors are important for getting better in my class”), affective-motivational adaptivity of error reactions (6 items; “When I say something wrong in my class, the class is still just as fun for me as always”), and action adaptivity of error reactions (7 items, e.g., “When something is too hard for me in my class, then it’s clear that I need to prepare better for class”).

#### Study strategies

We measured students’ study strategies using a measure originally developed by Morehead, Rhodes, and DeLozier (2016) and adapted by Rodriguez et al. (2018) for use with undergraduates. Students’ use of study strategies is measured by asking students to select the top three strategies they use from a provided list: “Absorbing lots of information the night before a test; condensing/Summarizing your notes; Make diagrams, charts, or pictures; Recopy your notes from memory; Recopy your notes word-for-word; Reread chapters, articles, notes, etc.; Study with friends; Test yourself with questions or practice problems; Underlining or highlighting while reading; Use flashcards; Watch/listen to recorded lessons either by instructor or from outside source (Khan Academy, YouTube, etc.); Other: write in.” Then, a separate item asks students to select which better describes their study patterns, “I most often space out my study sessions over multiple days/weeks” or “I most often do my studying right before the test.” We classified students’ study pattern as spacing or cramming based on their responses to these two questions. Students are classified as using spacing if they selected the spacing option to the second item and did **not** select “Absorbing lots of information the night before a test” as one of their top three study strategies in the prior item.

We had relatively low rates of endorsement for many of the study strategies. Therefore, we created an indicator variable to indicate whether a participant selected that they used any highly effective study strategy. Based on prior literature on effective study strategies, we identified self-testing, re-writing notes from memory, and creating visuals as the highly effective study strategies in the provided list. Self-testing and re-writing notes from memory were selected as highly effective strategies based on extensive evidence on the effectiveness of recall (Carpenter et al., 2008; Dunlosky et al., 2013). We also included creating visuals based on empirical studies have that have demonstrated the learning benefits of creating visuals (e.g., Ainsworth et al., 2011; Chularut & DeBacker, 2004; Fiorella & Mayer, 2016; Fiorella & Kuhlmann, 2020; Heideman et al., 2017; Schwamborn et al., 2010). Additionally, in the revised

Bloom’s taxonomy, creating is the highest-level activity (Krathwohl, 2002). Thus, we reasoned that a strategy involving a high level of cognitive engagement would be effective at promoting learning. Therefore, a participant was classified as using highly effective study strategies if they selected any of these three study strategies as one of their top three study strategies.

### Data analysis methods

Analyses were conducted in Mplus version 8 (Muthén & Muthén, 2017). First, a measurement model was estimated to confirm the factor structure of the eight latent variables: (1) fixed mindset, (2) growth mindset, (3) universal beliefs, (4) non-universal beliefs, (5) brilliance beliefs, (6) error beliefs, (7) affective-motivational adaptivity to errors, and (8) action adaptivity to errors. The measurement model also included two indicator variables, (1) spacing study sessions and (2) using highly effective study strategies. Both indicators were coded as Yes = 1 and No = 0. From this measurement model, bivariate correlations were derived to preliminarily assess the associations among all variables.

Following this, the hypothesized mediation model (Figure 1) was tested via a fully latent structural equation model (SEM) using Robust Maximum Likelihood (MLR) estimator. MLR adjusts the model fit indices to account for non-normality, which is commonly found within data using response scales of agreement, and was observed within the present data (Li, 2016; Zhong & Yuan, 2011). Covariances among error beliefs, action adaptivity, and motivational-affective adaptivity were freely estimated. To evaluate the fit of the data to the model, we examined several goodness-of-fit indices, specifically, the chi-square statistic, Comparative Fit Index (CFI), Tucker-Lewis Index (TLI), Root Mean Square Error of Approximation (RMSEA), Standardized Root Mean Square Residual (SRMR), and Chi-square/Degrees of Freedom (DF) ratio. Good model fit is typically indicated by CFI and TLI values close to or above 0.90, RMSEA below 0.06, SRMR below 0.08, and Chi-square/DF ratio of less than 2 (Hu & Bentler, 1999). Indirect effects were assessed using percentile bootstrap confidence intervals by recalculating the model with 1,000 resamples. In so doing, the indirect effect is based on the distribution of the resampled estimates versus a single product value, and thus, is likely to be more reliable than alternative estimates (Tibbe & Montoya, 2022).

## RESULTS

### Descriptive statistics

More than half of our participants (208/345; 60%) indicated they use at least one highly effective study strategy: self-testing, recopying notes from memory, or creating visuals. The most common study strategy reported by our participants was self-testing (167/345; 48%), followed by re-reading material (141/345; 41%), condensing/summarizing notes (131/345; 38%), and watching lectures and videos (125/345; 36%). The least commonly used strategies were recopying notes from memory (26/345; 7.5%), recopying notes word-for-word (47/345; 14%), creating visuals (47/345; 14%), underlining and highlighting while reading (49/345; 14%), and using flashcards (97/345; 28%). A minority of our participants (119/338; 35%) indicated that they space their study sessions out across time.

### Preliminary Results – Measurement Model

The measurement model fit the data adequately: χ^2^(902) = 1,517.95, p < .001, ^2^/df = 1.68; RMSEA = 0.044, 90% CI [0.041, 0.048]; CFI = 0.91; TLI = 0.90; SRMR = 0.05, with all item loadings at or above 0.48. This provides evidence of validity based on internal structure, suggesting that our measures functioned as expected in our sample. We assessed reliability by estimating McDonald’s Omega, which describes the proportion of the covariances among the indicators accounted for by the latent variable (McDonald., 1999). The omega of each measure used in this study was > 0.6, demonstrating acceptable reliability (Table 2; Hayes & Coutts, 2020).

**Table 2.**
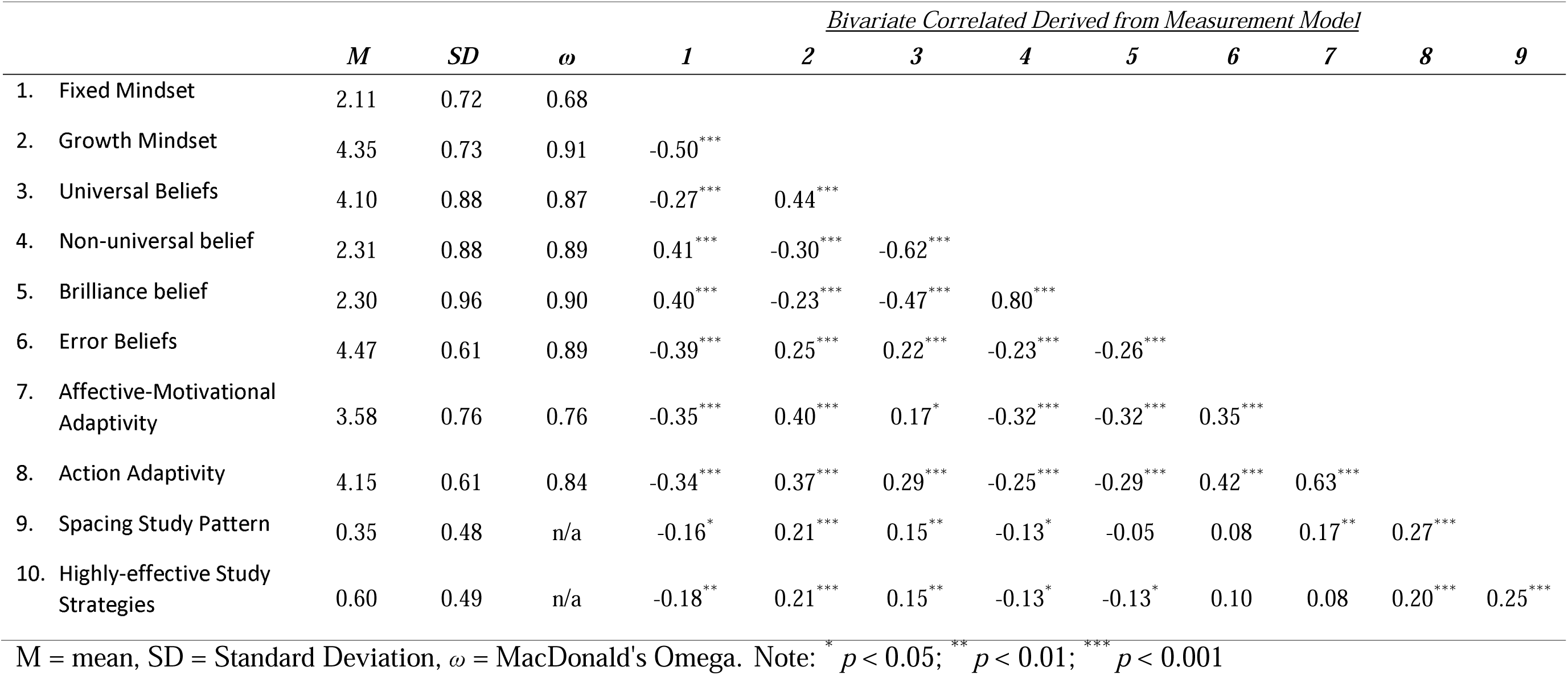
Descriptive statistics and bivariate correlations.

| | *M* | *SD* | $\omega$ | *1* | <u>*Bivariate Correlated Derived from Measurement Model*</u> | | | | | | | |
| --- | --- | --- | --- | --- | --- | --- | --- | --- | --- | --- | --- | --- |
|  |  |  |  |  | *2* | *3* | *4* | *5* | *6* | *7* | *8* | *9* |
| 1. Fixed Mindset | 2.11 | 0.72 | 0.68 |  |  |  |  |  |  |  |  |  |
| 2. Growth Mindset | 4.35 | 0.73 | 0.91 | -0.50*** |  |  |  |  |  |  |  |  |
| 3. Universal Beliefs | 4.10 | 0.88 | 0.87 | -0.27*** | 0.44*** |  |  |  |  |  |  |  |
| 4. Non-universal belief | 2.31 | 0.88 | 0.89 | 0.41*** | -0.30*** | -0.62*** |  |  |  |  |  |  |
| 5. Brilliance belief | 2.30 | 0.96 | 0.90 | 0.40*** | -0.23*** | -0.47*** | 0.80*** |  |  |  |  |  |
| 6. Error Beliefs | 4.47 | 0.61 | 0.89 | -0.39*** | 0.25*** | 0.22*** | -0.23*** | -0.26*** |  |  |  |  |
| 7. Affective-Motivational Adaptivity | 3.58 | 0.76 | 0.76 | -0.35*** | 0.40*** | 0.17* | -0.32*** | -0.32*** | 0.35*** |  |  |  |
| 8. Action Adaptivity | 4.15 | 0.61 | 0.84 | -0.34*** | 0.37*** | 0.29*** | -0.25*** | -0.29*** | 0.42*** | 0.63*** |  |  |
| 9. Spacing Study Pattern | 0.35 | 0.48 | n/a | -0.16* | 0.21*** | 0.15** | -0.13* | -0.05 | 0.08 | 0.17** | 0.27*** |  |
| 10. Highly-effective Study Strategies | 0.60 | 0.49 | n/a | -0.18** | 0.21*** | 0.15** | -0.13* | -0.13* | 0.10 | 0.08 | 0.20*** | 0.25*** |

M = mean, SD = Standard Deviation, $\omega$ = MacDonald's Omega. Note: \* $p < 0.05$ ; \*\* $p < 0.01$ ; \*\*\* $p < 0.001$

Table 2 presents the bivariate correlations from the measurement model. The associations were largely in the expected directions. All five lay theories correlated in the predicted directions with beliefs about errors and both types of reactions to errors. Additionally, fixed and growth mindset both significantly correlated in the expected directions with the use of both spacing study patterns and highly effective study strategies. Action adaptivity was positively associated with both spacing study patterns and using highly effective study strategies. Interestingly, affective-motivational adaptivity was only positively associated with spacing study patterns.

Counter to our expectations, beliefs about errors were not associated with either the use of spacing study patterns or high utility study strategies.

### Primary Analysis – Mediational Model

Our hypothesized model fit the data adequately: χ^2^(912) = 1,535.90, p < .001, ^2^/df = 1.68; RMSEA = 0.045, 90% CI [0.041, 0.048]; CFI = 0.91; TLI = 0.90; SRMR = 0.05.

Significant parameter estimates are presented in Figure 2 and all parameter estimates are available in Supplemental Material Section 2. As we hypothesized, fixed mindset was negatively associated with error beliefs and growth mindset was positively associated with both action and affective-motivational adaptivity. However, contrary to our hypothesis, universality, non-universality, and brilliance beliefs were not associated with error beliefs or adaptivity, either action or affective-motivational.

**Figure 2:**
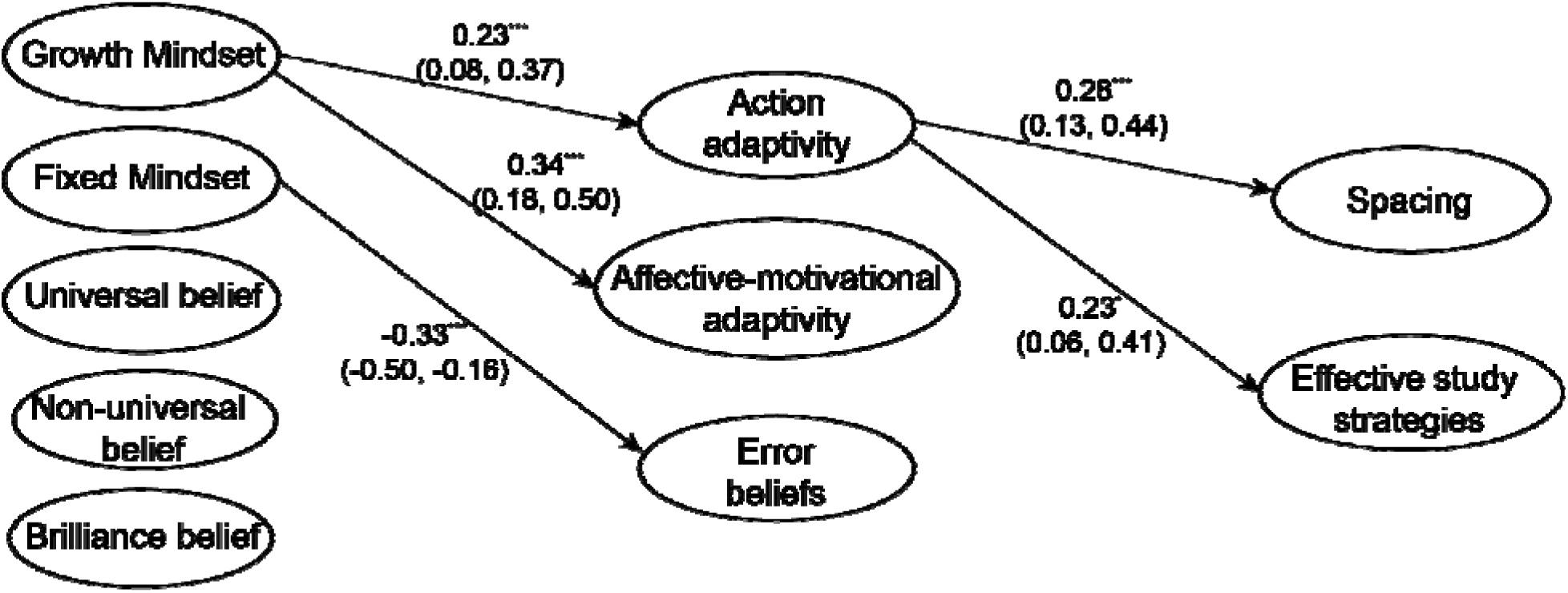
Structural Equation Modeling results showing standardized parameter estimates and 95% confidence intervals in parentheses. For simplicity, covariances and item loadings, as well as parameter estimates values non-significant at the p < 0.05 level are not shown. See full results in Supplemental Material section 2. Note: * *p* < 0.05; ** *p* < 0.01; *** *p* < 0.001 In alignment with our hypothesized model, action adaptivity was significantly positively associated with spacing out study sessions and using effective study strategies. However, contrary to our relationship, neither affective-motivational adaptivity nor error beliefs related to spacing or using highly effective study strategies.

We evaluated the indirect effects between growth mindset and spacing and using effective study strategies via traditional hypothesis testing as well as percentile bootstrap confidence intervals. We found that growth mindset significantly contributed to students’ spacing their study sessions through their action adaptivity toward errors (β = 0.06, 95% CI[0.02, 0.13], *p* = 0.028). Growth mindset contributed to students’ use of highly-effective study strategies through their action adaptivity toward errors (β = 0.05, 95% CI[0.01, 0.12], *p* = 0.073). Importantly, due to the non-significant p-value for the second indirect effect, caution is merited in interpreting this finding.

## DISCUSSION

Our results illuminate factors that relate to students’ use of effective study strategies. We found that growth mindset indirectly related to students’ study patterns, and that this relationship is mediated by their adaptivity to errors. However, contrary to our hypothesis, universality and brilliance beliefs did not relate to students’ beliefs about errors, reactions to errors, or study strategies, and students’ beliefs about errors and affective-motivational adaptivity did not relate to their study patterns or strategies. Our results imply that students who view abilities as improvable are more resilient to making errors while studying and thus are more likely to use more effective study strategies (*i.e.*, spacing out study sessions and self-testing, creating visuals, and writing notes from memory).

This study contributes to the growing literature base exploring mechanisms through which beliefs about abilities influence students’ academic outcomes. There is extensive prior research on the mechanisms through which beliefs about the malleability of intelligence (mindset) influence academic outcomes. These mechanisms include self-regulatory processes such as goal setting, operating, and monitoring (Burnette et al., 2013; Yan & Wang, 2021), and willingness to accept feedback (e.g., Sun & Huang, 2023). Our results add to growing evidence for a new mechanism: students’ reactions to errors and study strategies. Our results suggest that students who believe their intellectual abilities can improve are more likely to respond adaptively to making errors in the learning process and are more likely to space their study sessions out and use highly effective, error-prone study strategies. This finding corroborates prior work by Yan and Wang (2021) which found that intelligence mindset was indirectly related to studying behaviors, linked by goal orientation, which positively related to both growth mindset and use of effective study strategies. Our findings also build on prior work by Ng (2018), which provided neuroscientific evidence that individuals with stronger growth mindset pay more attention to errors and learn more from them.

Our study also explored possible effects of beliefs about distribution of potential (universality beliefs) and whether high intellectual ability is required for success (brilliance belief) on study strategies. Contrary to our expectations, we found that neither universality nor brilliance were related to beliefs about and reactions to errors as well as study patterns and strategies. Our results indicate that some beliefs about abilities relate to study strategy use, but not others. This corroborates prior research indicating that mindset, universality, and brilliance uniquely relate to different psychosocial and academic outcomes (Limeri et al., 2023). Our results provide further evidence that these beliefs have different modes of action (mediators) through which they impact students’ outcomes. Future work should continue to explore which mediators and outcomes are influenced by each of the three lay theories about abilities and explore the mechanisms underlying these differential connections.

### Implications

These results may potentially be used to inform the design of study strategy interventions to improve students’ academic success by encouraging students to respond adaptively to making errors during their learning. Interventions targeting study strategies have had mixed success (McDaniel & Einstein, 2020). Research has found that educating students about the effectiveness of strategies is by itself insufficient to motivate behavioral change (Biwer et al., 2020; Dye & Stanton, 2017; Morehead et al., 2016; Rea et al., 2022; Wang, Muenks, & Yan, 2023). McDaniel and Einstein (2020) conceptualized the knowledge, belief, commitment, and planning (KBCP) framework, which posits that knowledge is only a first step, but students must also be persuaded that effective study strategies will work better, commit to using effective strategies, and create a feasible plan for using effective strategies (McDaniel & Einstein, 2020). Our results suggest that helping students respond positively to making errors may be another important part of helping them adopt effective learning strategies. Incorporating elements about responding adaptively to making errors while learning and growth mindset may increase the efficacy of interventions. It will be important to test if incorporating these elements into interventions will increase their success.

Our results suggest that future studies investigating student learning and study strategies should consider students’ views of their abilities and their adaptive reactions to errors. Rea et al. (2022) found that despite recognizing the benefits of more effective study strategies like pretesting and interleaving, participants reported relying primarily on passive strategies. Rea et al. (2022) identified barriers to adopting effective study strategies, including perceptions that they are too time-consuming, require excessive effort, or having low self-efficacy for using these strategies well. They also identified barriers to effective metacognitive strategies (i.e., goal setting, help-seeking, study planning, and focus of time and effort), the primary barrier being psychological cost (i.e., these strategies would induce nervousness and anxiety). Future research should explore whether helping students develop positive beliefs about and reactions to errors may reduce some of these perceived barriers or increase students’ confidence that making errors is an indicator that they are studying effectively. Our results suggest that simultaneously targeting growth mindset beliefs may prove fruitful since they relate to students’ error adaptivity. Future work could develop and implement interventions that target study strategies, the positive role of errors in studying, as well as promote a growth mindset. This is promising because prior work indicates that mindsets are malleable, even for undergraduates (Limeri et al., 2020).

Helping students shift away from less efficient strategies towards more efficient study strategies can benefit all students. We propose it is possible that these efforts may yield disproportionate benefits for students who face the most barriers in their educational journeys. While more privileged students may be able to compensate for using inefficient study strategies by increasing the total amount of time they spend studying, this option may not be available for students who face constraints, such as working to support themselves and their family, or those with caretaking responsibilities. Thus, students who are low-income and/or are caretakers for family may have the least time available for studying and thus may stand the most to gain from using their time as efficiently as possible. There is some evidence that study strategies may be particularly important for students who face additional barriers in their education. DiBenedetto (2010) surveyed 248 undergraduates and found that there was a stronger relationship between self-regulated learning strategies and college GPA for first-generation students than for continuing-generation students.

There is some promising evidence that metacognitive interventions that teach students about effective study strategies have potential to improve equity in student achievement (Muteti et al., 2023; Rodriguez et al., 2018). Muteti and colleagues (2023) found that first-generation students reported using self-testing and practice problems less often than continuing generation students, but that a 50-minute lesson about metacognition reduced this gap. Rodriguez and colleagues (2018) found that the performance gap between white students and Persons Excluded due to Ethnicity or Race (PEERs) was higher when PEERs did not self-test, and reduced when PEERs engage in self-testing. They also found that their light-touch intervention was successful in encouraging all students, including PEERs, to engage in self-testing more frequently (Rodriguez et al., 2018).

### Limitations and Future Directions

Our study is cross-sectional and as such we are unable to draw conclusions about causality among the variables studied. We hypothesize a mediation model based on theoretical linkages. For example, although we hypothesize that students’ mindset beliefs influence their reactions to errors, it is possible that the reverse is true. Perhaps frequently engaging in self-testing and making errors teaches students that their abilities can improve and consequently strengthens their growth mindset beliefs. Future work could improve on this by collecting data longitudinally to investigate causal relationships. Alternatively, future work could take a qualitative approach to gain more insight into students’ reasoning for choosing their study strategies and the role that errors play in their choices about studying.

Our study was conducted in a biology class at a single, primarily white university located in the United States. Thus, our sample is limited in demographic diversity. Caution should be taken in generalizing these findings to other student populations, including students in other STEM disciplines, at different institution types (e.g., teaching-focused, community colleges, minority-serving), with different racial/ethnic identities, and in different countries. For example, previous studies have indicated that study strategy use differs across disciplines (Chouvalova et al., 2022; Lawson, 1979). Engineering students placed greater emphasis (relative to biology students) on the importance of self-correction, and making multiple approaches and attempts (Chouvalova et al., 2022). It is possible that epistemic differences and differences in types of problems typically encountered between fields relate to different beliefs about abilities and approaches to studying. Future work should examine how the patterns identified here may generalize to other populations and explore how these variables and processes interact with students’ intersectional identities.

An important limitation is that we dichotomized study strategies as either highly effective or not. An underlying assumption that we are making is that students who are using study strategies traditionally identified as being ineffective, like using flashcards and rereading material, are indeed using them ineffectively. Cognitive processes occurring during passive study strategies that are traditionally categorized as ineffective, such as using flashcards and rereading material, may involve higher orders of cognition like making connections and retrieving prior knowledge. For example, flashcards could be used to engage in self-testing if the student is actively recalling information before reviewing the other side of the card, or as passive re-reading if students are flipping to the other side without engaging in difficult recall (Miyatsu et al., 2018). If students use flashcards strategically, they can be an effective study strategy (Kornell et al., 2009; Miyatsu et al., 2018; Schmidmaier et al., 2011; Senzaki et al., 2017). Miyatsu et al. (2018) reviewed literature and identified some ways students can implement flashcards effectively, such as retaining cards even after successfully recalling them and spacing out sessions. Kornell (2009) conducted a lab experiment with undergraduates at a research-intensive public institution in the Western United States on the spacing of flashcard sessions and found that studying large stacks of flashcards was more effective than separating cards into smaller piles because it created more spacing between practice sessions for each word. The utility of flash cards also depends on the learning goal; for example, they are particularly effective for rote memorization of definitions of terms (Miyatsu et al., 2018). We took a conservative approach of classifying study strategies that could be used ineffectively as not highly effective, since their use is not a reliable sign of using effective study strategies. With flash card use in particular, studies suggest that most of the time, students will stop practicing a flashcard if it is successfully retrieved once (Miyatsu et al., 2018) and typically, students resort to flashcards for lower-order processes like memorizing vocabulary, instead of gaining a detailed understanding of a concept or knowledge application. In fact, Hartwig and Dunlosky (2012) concluded that flashcard use is not related to students’ GPA.

Likewise, we are also assuming that students who report using study strategies identified as being effective, like retrieval and spacing, are indeed using them effectively. However, with effective study strategies, certain nuances may undermine their effectiveness. With retrieval testing, Wooldridge et al. (2014) found that this method is only useful for questions that are repeated on the practice test and actual test but not for topically related items, which indicates that perhaps retrieval testing is not as effective for transferring knowledge. For example, a student using retrieval testing may be memorizing the questions and solutions and in doing so, engaging in lower orders of cognition. Future studies could investigate how students implement their study strategies and the factors that relate to the quality of implementation of effective strategies.

## CONCLUSION

We find partial support for our hypothesized mediation model (Figure 1). In alignment with our hypothesis, we find that students’ growth mindset beliefs are positively associated with reacting adaptively to making errors, which is in turn positively associated with spacing study sessions and using highly effective study strategies. In contrast, we did not find support for relationships between other types of beliefs about abilities (universality and brilliance beliefs) and reactions to errors or study strategies These results build on the literature base exploring the mechanisms through which students’ mindset beliefs influence their academic outcomes. Our results also add new knowledge about the motivational factors that relate to students’ choices about how and when to study. This knowledge could be used to inform the design of future study strategy interventions, focusing not just on informing students about the effectiveness of strategies, but motivating them to adapt positively to errors and enable them to adopt effective strategies more readily.

## LIST OF ABBREVIATIONS

CFI: Comparative Fit Index
DF: Degrees of Freedom
MLR: Robust Maximum Likelihood
RMSEA: Root Mean Squared Error of Approximation
SD: Standard Deviation
SEM: Structural Equation Model
SRMR: Standardized Root Mean Squared Residual
STEM: Science, technology, engineering, and mathematics
TLI: Tucker-Lewis Index
ULTrA: Undergraduate Lay Theories of Abilities Survey

## DECLARATIONS

### Availability of data and materials

The dataset supporting the conclusions of this article is included within the article and its additional files.

### Competing interests

The authors declare that they have no competing interests.

### Funding

This work was supported by start-up funds from Texas Tech University (TTU) and student support was provided by The Center for Transformative Undergraduate Experiences at TTU. Funders had no role in the conceptualization, design, data collection, analysis, decision to publish, or preparation of the manuscript.

### Authors’ contributions

AC contributed to data analysis and interpretation and drafting the manuscript. AN contributed to data analysis and interpretation and drafting the manuscript. DJM contributed to data analysis and interpretation and drafting the manuscript. MA contributed to conceptualizing and designing the study and data collection. SD contributed to conceptualizing and designing the study and data collection. FDA contributed to conceptualizing and designing the study and data collection. LBL contributed to conceptualizing and designing the study, data collection, analysis, and interpretation, and drafting and substantially revising the writing. All authors read and approved the final manuscript.

## Acknowledgements

We would like to thank the instructor who taught the course for assisting with disseminating the study and providing extra credit as incentive.

## Notes

### Competing Interest Statement

The authors have declared no competing interest.

### Summary of Updates

The revision reflects changes in response to reviewer feedback.

