## Supplemental Material for "Undergraduates’ reactions to errors mediates the association between growth mindset and study strategies"

**Index**

Section 1: Data collection measures, pgs S2-S6

Section 2: MPLUS input and Full SEM results, pgs S7-S31

**Supplemental Material Section 1: Data Collection Measures**

**ULTrA Survey:** mindset, universality, brilliance beliefs (Limeri et al., 2023)

Limeri, L. B., Carter, N. T., Lyra, F., Martin, J., Mastronardo, H., Patel, J., & Dolan, E. L. (2023). Undergraduate Lay Theories of Abilities: Mindset, universality, and brilliance beliefs uniquely predict undergraduate educational outcomes. *CBE – Life Sciences Education, 22*(4), ar40. <https://doi.org/10.1187/cbe.22-12-0250>

Instructions: “Please indicate the extent to which you agree or disagree with the following statements. There are no correct answers, we want to understand how you think about these ideas. Note that STEM stands for Science, Technology, Engineering, and Mathematics. STEM professionals are individuals in a career in a STEM field, such as scientists, engineers, medical doctors, and other healthcare professionals.”

Response Scale: 1 = “strongly disagree”; 2 = “Somewhat disagree”; 3 = “Neither agree nor disagree”; 4 = “Somewhat agree”’ 5 = “Strongly agree”; “Prefer not to respond”

Fixed mindset (5 items)

- At the end of college, my ability to analyze information will be at about the same level that it is now.
- How well I learn is something that I cannot change very much.
- My ability to apply knowledge will change very little over time.
- I will never be able to reach the highest level of intellectual ability.
- It would be very difficult for me to improve how well I can apply knowledge.

Growth mindset (5 items)

- I can become as good at analyzing information as highly successful STEM professionals if I try hard enough.
- If I want to, I can become as effective at applying knowledge as STEM experts.
- I can become excellent at applying knowledge to solve challenging problems.
- If I try, I can become as effective at learning as STEM experts.
- I could improve my intellectual abilities to the same level as successful STEM professionals.

Non-universal belief (5 items)

- Even if they try, some people could never become as effective at analyzing information as their peers.
- Only people with a natural talent can become good enough at applying knowledge to solve the most difficult problems.
- Only people with a natural talent can become excellent at analyzing information.
- Some people will always be less effective at learning than those who have a natural talent for it.
- Only some people have the intellectual ability to become a successful STEM professional.

Universal belief (5 items)

- With enough hard work, anyone could become as good at analyzing information as highly successful STEM professionals.
- Anyone could become as effective at learning as highly successful STEM students.
- Anyone who tries could become as good at applying knowledge as STEM experts.
- Everyone has the intellectual ability to become a successful STEM professional if they want to.
- With enough motivation, anyone can become as good at applying knowledge as high achieving STEM students.

Brilliance belief (5 items)

- Excelling in STEM requires natural talent.
- People who are highly successful in STEM have a natural talent for it.
- Becoming a top student in STEM requires an innate talent that just can't be taught.
- People have to be naturally talented to excel in challenging STEM courses.
- Being a highly successful STEM professional requires natural talent that just can't be taught.

**Beliefs about errors** (Tulis et al., 2017)

Tulis, M., Steuer, G., & Dresel, M. (2017). Positive beliefs about errors as an important element of adaptive individual dealing with errors during academic learning. *Educational Psychology*, 1–20. <https://doi.org/10.1080/01443410.2017.1384536>

Instructions: “Please indicate the extent to which you agree or disagree with the following statements. There are no correct answers, we want to understand how you think about these ideas.”

Response Scale: 1 = “strongly disagree”; 2 = “Somewhat disagree”; 3 = “Neither agree nor disagree”; 4 = “Somewhat agree”’ 5 = “Strongly agree”; “Prefer not to respond”

- I can learn something from my errors in my class.
- Errors are important for getting better in my class.
- I develop new skills by making errors in my class.
- My errors help me to improve my skills in my class.
- Errors in my class help me to be better at it later on.

**Error Adaptivity** (Tulis et al., 2017)

Instructions: “Please indicate the extent to which you agree or disagree with the following statements. There are no correct answers, we want to understand how you think about these ideas.”

Response Scale: 1 = “strongly disagree”; 2 = “Somewhat disagree”; 3 = “Neither agree nor disagree”; 4 = “Somewhat agree”’ 5 = “Strongly agree”; “Prefer not to respond”

Affective-motivational adaptivity (6 items)

- When I say something wrong in my class, then the class is ruined as far as I am concerned. (reverse-scored)
- When I say something wrong in my class, the class is still just as fun for me as always.
- When I can't do something in my class, the lessons in the future will still be just as fun for me as always.
- When I can't solve a problem in my class, then I have less motivation next time around. (reverse-scored)
- When I make an error in my class, then I will have less fun in my class later on. (reverse-scored)
- When I can't do something in my class, I still want to keep working on it.

Action adaptivity (7 items)

- When I can’t do something in my class, then I try even harder the next time around.
- When something is too hard for me in my class, then it's clear that I need to prepare better for class.
- When I make an error in my class, then I set a goal to try to improve myself.
- When I make a mistake in my class, then I know where I will have to focus my efforts next time around.
- When I do something wrong in my class, then I specifically try to work it out.
- When I am not able to solve a problem in my class, this helps me to know where I can improve myself.
- When I can't solve a problem in my class, then I practice these types of exercises on my own.

**Study Strategies** (Rodriguez et al., 2018)

original measure from

Morehead K, Rhodes MG, & DeLozier S (2016) Instructor and student knowledge of study strategies. Memory 24(2): 257–271.

Current version was adapted by

Rodriguez, F., Rivas, M. J., Matsumura, L. H., Warschauer, M., & Sato, B. K. (2018). How do students study in STEM courses? Findings from a light-touch intervention and its relevance for underrepresented students. PLOS ONE, 13(7), e0200767.

Select the top three study strategies you use most regularly. Please select ONLY 3.

- Absorbing lots of information the night before a test
- Condensing/Summarizing your notes
- Make diagrams, charts, or pictures
- Recopy your notes from memory
- Recopy your notes word-for-word
- Reread chapters, articles, notes, etc.
- Study with friends
- Test yourself with questions or practice problems
- Underlining or highlighting while reading
- Use flashcards
- Watch/listen to recorded lessons either by instructor or from outside source (Khan Academy, YouTube, etc.)
- Other: write in

Which of the following best describes your study patterns?

- I most often space out my study sessions over multiple days/weeks
- I most often do my studying right before the test

When studying, how do you generally decide what class to study for first?

- Whatever's due soonest/overdue
- Whatever I haven't studied for the longest time
- Whatever I find interesting
- Whatever I feel I'm doing the worst in
- I plan my study schedule ahead of time and I study whatever I've scheduled
- Other: __________

**Attention check questions:**

- This is a control question. Please select "Strongly disagree."
- This is a control question. Please select “Prefer not to respond.”

**Demographic questions**

What is your current GPA (on 4.0 scale)?,

Which of the following accurately describes your family's education?

- Continuing generation: At least one of my parent(s)/guardian(s) has earned a 4-year college degree.
- First generation: None of my parent(s)/guardian(s) has earned a 4-year college degree.
- Prefer not to respond.

With which race(s) and ethnicity/ies do you identify? Select all that apply:

- African American or Black
- East Asian (e.g., China and Japan)
- South Asian (e.g., the Indian sub-continent)
- Southeast Asian (e.g., Vietnam)
- Latinx or Hispanic
- Middle Eastern or North African
- Native American or Alaskan Native
- Native Hawaiian or Pacific Islander
- White
- Other: ____
- Prefer not to respond

What gender do you identify as?

- Man
- Woman
- Non-binary
- Not listed above: _____
- Prefer not to respond

**Supplemental Material Section 2: MPLUS input and Full SEM results**

```
SEM with MLR Estimator

  ANALYSIS:
    	TYPE = GENERAL;
    	ESTIMATOR = mlr;

  MODEL:
!Fixed Mindset factor
      fxblf  by   fxblf1 fxblf2 fxblf3 fxblf4 fxblf5;
!Growth Mindset factor
      grblf  by   grblf1 grblf2 grblf3 grblf4 grblf5;
!Universal Beliefs factor
      unblf  by   unblf1 unblf2 unblf3 unblf4 unblf5;
!Non-Universal Beliefs factor
      nunblf by   nunblf1 nunblf2 nunblf3 nunblf4 nunblf5;
!Brilliance Beliefs factor
      brblf  by   brblf1 brblf2 brblf3 brblf4 brblf5;

!Error Beliefs factor
      errblf   by   errblf1 errblf2 errblf3 errblf4 errblf5;
!Affective-Motivational Adaptivity factor
      m_adap   by   m_adap1 m_adap2 m_adap3 m_adap4 m_adap5 m_adap6;
!Action Adaptivity factor
      a_adap   by   a_adap1 a_adap2 a_adap3 a_adap4 a_adap5 a_adap6 a_adap7;

!Regressions for Mediators
      errblf on fxblf grblf unblf nunblf brblf;

      m_adap on fxblf grblf unblf nunblf brblf;

      a_adap on fxblf grblf unblf nunblf brblf;

!Regressions for Outcomes
      space on errblf m_adap a_adap;
      high_utl on errblf m_adap a_adap;

!Correlated Errors/Disturbances
      errblf with m_adap a_adap;
      m_adap with a_adap;

Model indirect:
      space ind grblf;
      high_utl ind grblf;

      space ind fxblf;
      high_utl ind fxblf;

  OUTPUT:
  	SAMPSTAT;
  	CINTERVAL;
  	STDYX;
  	STANDARDIZED;
  	MODINDICES;
      	TECH4;

MODEL FIT INFORMATION

Number of Free Parameters                      168

Loglikelihood

          H0 Value                      -17512.842
          H0 Scaling Correction Factor      1.3429
            for MLR
          H1 Value                      -16645.108
          H1 Scaling Correction Factor      1.1631
            for MLR

Information Criteria

          Akaike (AIC)                   35361.685
          Bayesian (BIC)                 36007.400
          Sample-Size Adjusted BIC       35474.458
            (n* = (n + 2) / 24)

Chi-Square Test of Model Fit

          Value                           1535.902*
          Degrees of Freedom                   912
          P-Value                           0.0000
          Scaling Correction Factor         1.1299
            for MLR

*   The chi-square value for MLM, MLMV, MLR, ULSMV, WLSM and WLSMV cannot be used
    for chi-square difference testing in the regular way.  MLM, MLR and WLSM
    chi-square difference testing is described on the Mplus website.  MLMV, WLSMV,
    and ULSMV difference testing is done using the DIFFTEST option.

RMSEA (Root Mean Square Error Of Approximation)

          Estimate                           0.045
          90 Percent C.I.                    0.041  0.048
          Probability RMSEA <= .05           0.991

CFI/TLI

          CFI                                0.905
          TLI                                0.897

Chi-Square Test of Model Fit for the Baseline Model

          Value                           7540.363
          Degrees of Freedom                   990
          P-Value                           0.0000

SRMR (Standardized Root Mean Square Residual)

          Value                              0.056


STANDARDIZED MODEL RESULTS


STDYX Standardization

                                                    Two-Tailed
                    Estimate       S.E.  Est./S.E.    P-Value

 FXBLF    BY
    FXBLF1             0.478      0.061      7.801      0.000
    FXBLF2             0.572      0.055     10.403      0.000
    FXBLF3             0.504      0.060      8.415      0.000
    FXBLF4             0.505      0.062      8.106      0.000
    FXBLF5             0.678      0.058     11.718      0.000

 GRBLF    BY
    GRBLF1             0.743      0.038     19.345      0.000
    GRBLF2             0.819      0.028     29.280      0.000
    GRBLF3             0.774      0.036     21.771      0.000
    GRBLF4             0.880      0.020     43.092      0.000
    GRBLF5             0.860      0.025     34.089      0.000

 UNBLF    BY
    UNBLF1             0.821      0.032     25.438      0.000
    UNBLF2             0.831      0.028     29.501      0.000
    UNBLF3             0.854      0.027     31.543      0.000
    UNBLF4             0.786      0.029     27.240      0.000
    UNBLF5             0.860      0.024     35.657      0.000

 NUNBLF   BY
    NUNBLF1            0.630      0.044     14.470      0.000
    NUNBLF2            0.795      0.039     20.617      0.000
    NUNBLF3            0.756      0.052     14.483      0.000
    NUNBLF4            0.667      0.040     16.619      0.000
    NUNBLF5            0.697      0.038     18.442      0.000

 BRBLF    BY
    BRBLF1             0.732      0.033     21.937      0.000
    BRBLF2             0.647      0.037     17.500      0.000
    BRBLF3             0.864      0.021     40.907      0.000
    BRBLF4             0.855      0.023     37.168      0.000
    BRBLF5             0.863      0.020     43.455      0.000

 ERRBLF   BY
    ERRBLF1            0.594      0.043     13.916      0.000
    ERRBLF2            0.739      0.040     18.346      0.000
    ERRBLF3            0.824      0.035     23.278      0.000
    ERRBLF4            0.914      0.023     40.393      0.000
    ERRBLF5            0.853      0.029     29.361      0.000

 M_ADAP   BY
    M_ADAP1            0.572      0.049     11.598      0.000
    M_ADAP2            0.637      0.053     11.908      0.000
    M_ADAP3            0.577      0.056     10.224      0.000
    M_ADAP4            0.538      0.059      9.109      0.000
    M_ADAP5            0.630      0.063     10.062      0.000
    M_ADAP6            0.590      0.057     10.322      0.000

 A_ADAP   BY
    A_ADAP1            0.683      0.035     19.291      0.000
    A_ADAP2            0.516      0.045     11.340      0.000
    A_ADAP3            0.683      0.035     19.714      0.000
    A_ADAP4            0.701      0.044     15.829      0.000
    A_ADAP5            0.756      0.033     23.188      0.000
    A_ADAP6            0.599      0.045     13.215      0.000
    A_ADAP7            0.615      0.043     14.139      0.000

 ERRBLF   ON
    FXBLF             -0.327      0.086     -3.797      0.000
    GRBLF              0.032      0.079      0.402      0.688
    UNBLF              0.106      0.096      1.104      0.269
    NUNBLF             0.102      0.156      0.652      0.514
    BRBLF             -0.151      0.125     -1.207      0.227

 M_ADAP   ON
    FXBLF             -0.104      0.107     -0.974      0.330
    GRBLF              0.338      0.084      4.050      0.000
    UNBLF             -0.183      0.099     -1.849      0.064
    NUNBLF            -0.167      0.149     -1.122      0.262
    BRBLF             -0.148      0.128     -1.160      0.246

 A_ADAP   ON
    FXBLF             -0.164      0.096     -1.701      0.089
    GRBLF              0.226      0.075      3.022      0.003
    UNBLF              0.140      0.090      1.549      0.121
    NUNBLF             0.150      0.156      0.959      0.337
    BRBLF             -0.226      0.129     -1.756      0.079

 SPACE    ON
    ERRBLF            -0.050      0.057     -0.870      0.385
    M_ADAP             0.024      0.088      0.269      0.788
    A_ADAP             0.283      0.081      3.501      0.000

 HIGH_UTL ON
    ERRBLF             0.019      0.066      0.281      0.779
    M_ADAP            -0.051      0.092     -0.550      0.582
    A_ADAP             0.231      0.089      2.585      0.010

 ERRBLF   WITH
    M_ADAP             0.238      0.072      3.328      0.001
    A_ADAP             0.305      0.067      4.585      0.000

 M_ADAP   WITH
    A_ADAP             0.557      0.069      8.129      0.000

 GRBLF    WITH
    FXBLF             -0.497      0.062     -8.018      0.000

 UNBLF    WITH
    FXBLF             -0.274      0.072     -3.794      0.000
    GRBLF              0.435      0.065      6.719      0.000

 NUNBLF   WITH
    FXBLF              0.405      0.074      5.456      0.000
    GRBLF             -0.297      0.063     -4.730      0.000
    UNBLF             -0.621      0.054    -11.474      0.000

 BRBLF    WITH
    FXBLF              0.405      0.061      6.601      0.000
    GRBLF             -0.231      0.055     -4.239      0.000
    UNBLF             -0.470      0.053     -8.927      0.000
    NUNBLF             0.803      0.033     24.001      0.000

 HIGH_UTL WITH
    SPACE              0.208      0.052      4.005      0.000

 Intercepts
    FXBLF1             1.877      0.059     32.052      0.000
    FXBLF2             2.134      0.067     31.878      0.000
    FXBLF3             1.850      0.057     32.230      0.000
    FXBLF4             1.801      0.049     37.027      0.000
    FXBLF5             2.148      0.075     28.759      0.000
    GRBLF1             5.062      0.298     16.985      0.000
    GRBLF2             4.867      0.300     16.200      0.000
    GRBLF3             6.052      0.435     13.907      0.000
    GRBLF4             5.121      0.280     18.294      0.000
    GRBLF5             4.786      0.280     17.096      0.000
    NUNBLF1            2.223      0.071     31.216      0.000
    NUNBLF2            1.820      0.052     35.118      0.000
    NUNBLF3            1.900      0.071     26.680      0.000
    NUNBLF4            2.171      0.072     30.073      0.000
    NUNBLF5            1.905      0.053     36.044      0.000
    UNBLF1             4.504      0.256     17.566      0.000
    UNBLF2             4.188      0.226     18.560      0.000
    UNBLF3             4.181      0.223     18.726      0.000
    UNBLF4             3.277      0.165     19.879      0.000
    UNBLF5             4.247      0.248     17.110      0.000
    BRBLF1             2.103      0.062     33.733      0.000
    BRBLF2             2.440      0.087     28.019      0.000
    BRBLF3             1.851      0.050     36.968      0.000
    BRBLF4             1.885      0.054     35.165      0.000
    BRBLF5             1.784      0.049     36.614      0.000
    ERRBLF1            8.070      0.495     16.302      0.000
    ERRBLF2            5.937      0.387     15.332      0.000
    ERRBLF3            5.676      0.349     16.249      0.000
    ERRBLF4            5.678      0.400     14.199      0.000
    ERRBLF5            6.024      0.437     13.800      0.000
    M_ADAP1            3.616      0.166     21.787      0.000
    M_ADAP2            3.054      0.119     25.728      0.000
    M_ADAP3            3.055      0.113     26.953      0.000
    M_ADAP4            2.462      0.078     31.620      0.000
    M_ADAP5            3.150      0.125     25.204      0.000
    M_ADAP6            3.955      0.206     19.206      0.000
    A_ADAP1            4.453      0.196     22.725      0.000
    A_ADAP2            5.701      0.312     18.252      0.000
    A_ADAP3            4.688      0.220     21.277      0.000
    A_ADAP4            6.019      0.358     16.833      0.000
    A_ADAP5            4.981      0.282     17.671      0.000
    A_ADAP6            5.652      0.363     15.562      0.000
    A_ADAP7            3.295      0.140     23.587      0.000
    SPACE              0.740      0.042     17.615      0.000
    HIGH_UTL           1.232      0.068     18.177      0.000

 Variances
    FXBLF              1.000      0.000    999.000    999.000
    GRBLF              1.000      0.000    999.000    999.000
    UNBLF              1.000      0.000    999.000    999.000
    NUNBLF             1.000      0.000    999.000    999.000
    BRBLF              1.000      0.000    999.000    999.000

 Residual Variances
    FXBLF1             0.772      0.059     13.184      0.000
    FXBLF2             0.673      0.063     10.687      0.000
    FXBLF3             0.746      0.060     12.354      0.000
    FXBLF4             0.745      0.063     11.822      0.000
    FXBLF5             0.541      0.078      6.899      0.000
    GRBLF1             0.448      0.057      7.846      0.000
    GRBLF2             0.329      0.046      7.166      0.000
    GRBLF3             0.400      0.055      7.270      0.000
    GRBLF4             0.225      0.036      6.254      0.000
    GRBLF5             0.260      0.043      5.987      0.000
    NUNBLF1            0.603      0.055     10.996      0.000
    NUNBLF2            0.368      0.061      5.993      0.000
    NUNBLF3            0.428      0.079      5.416      0.000
    NUNBLF4            0.555      0.054     10.350      0.000
    NUNBLF5            0.514      0.053      9.748      0.000
    UNBLF1             0.327      0.053      6.166      0.000
    UNBLF2             0.309      0.047      6.586      0.000
    UNBLF3             0.271      0.046      5.869      0.000
    UNBLF4             0.382      0.045      8.426      0.000
    UNBLF5             0.260      0.041      6.275      0.000
    BRBLF1             0.465      0.049      9.526      0.000
    BRBLF2             0.581      0.048     12.138      0.000
    BRBLF3             0.254      0.036      6.948      0.000
    BRBLF4             0.269      0.039      6.828      0.000
    BRBLF5             0.256      0.034      7.459      0.000
    ERRBLF1            0.647      0.051     12.777      0.000
    ERRBLF2            0.454      0.060      7.613      0.000
    ERRBLF3            0.320      0.058      5.489      0.000
    ERRBLF4            0.165      0.041      3.993      0.000
    ERRBLF5            0.272      0.050      5.473      0.000
    M_ADAP1            0.673      0.056     11.947      0.000
    M_ADAP2            0.594      0.068      8.721      0.000
    M_ADAP3            0.667      0.065     10.256      0.000
    M_ADAP4            0.711      0.063     11.199      0.000
    M_ADAP5            0.602      0.079      7.625      0.000
    M_ADAP6            0.652      0.067      9.663      0.000
    A_ADAP1            0.533      0.048     11.013      0.000
    A_ADAP2            0.734      0.047     15.654      0.000
    A_ADAP3            0.533      0.047     11.268      0.000
    A_ADAP4            0.508      0.062      8.177      0.000
    A_ADAP5            0.428      0.049      8.674      0.000
    A_ADAP6            0.641      0.054     11.810      0.000
    A_ADAP7            0.622      0.053     11.626      0.000
    SPACE              0.921      0.030     30.331      0.000
    HIGH_UTL           0.955      0.025     37.983      0.000
    ERRBLF             0.825      0.047     17.709      0.000
    M_ADAP             0.761      0.058     13.128      0.000
    A_ADAP             0.789      0.051     15.350      0.000


R-SQUARE

    Observed                                        Two-Tailed
    Variable        Estimate       S.E.  Est./S.E.    P-Value

    FXBLF1             0.228      0.059      3.900      0.000
    FXBLF2             0.327      0.063      5.201      0.000
    FXBLF3             0.254      0.060      4.208      0.000
    FXBLF4             0.255      0.063      4.053      0.000
    FXBLF5             0.459      0.078      5.859      0.000
    GRBLF1             0.552      0.057      9.673      0.000
    GRBLF2             0.671      0.046     14.640      0.000
    GRBLF3             0.600      0.055     10.885      0.000
    GRBLF4             0.775      0.036     21.546      0.000
    GRBLF5             0.740      0.043     17.045      0.000
    NUNBLF1            0.397      0.055      7.235      0.000
    NUNBLF2            0.632      0.061     10.308      0.000
    NUNBLF3            0.572      0.079      7.242      0.000
    NUNBLF4            0.445      0.054      8.310      0.000
    NUNBLF5            0.486      0.053      9.221      0.000
    UNBLF1             0.673      0.053     12.719      0.000
    UNBLF2             0.691      0.047     14.751      0.000
    UNBLF3             0.729      0.046     15.772      0.000
    UNBLF4             0.618      0.045     13.620      0.000
    UNBLF5             0.740      0.041     17.828      0.000
    BRBLF1             0.535      0.049     10.969      0.000
    BRBLF2             0.419      0.048      8.750      0.000
    BRBLF3             0.746      0.036     20.454      0.000
    BRBLF4             0.731      0.039     18.584      0.000
    BRBLF5             0.744      0.034     21.727      0.000
    ERRBLF1            0.353      0.051      6.958      0.000
    ERRBLF2            0.546      0.060      9.173      0.000
    ERRBLF3            0.680      0.058     11.639      0.000
    ERRBLF4            0.835      0.041     20.196      0.000
    ERRBLF5            0.728      0.050     14.680      0.000
    M_ADAP1            0.327      0.056      5.799      0.000
    M_ADAP2            0.406      0.068      5.954      0.000
    M_ADAP3            0.333      0.065      5.112      0.000
    M_ADAP4            0.289      0.063      4.554      0.000
    M_ADAP5            0.398      0.079      5.031      0.000
    M_ADAP6            0.348      0.067      5.161      0.000
    A_ADAP1            0.467      0.048      9.645      0.000
    A_ADAP2            0.266      0.047      5.670      0.000
    A_ADAP3            0.467      0.047      9.857      0.000
    A_ADAP4            0.492      0.062      7.915      0.000
    A_ADAP5            0.572      0.049     11.594      0.000
    A_ADAP6            0.359      0.054      6.608      0.000
    A_ADAP7            0.378      0.053      7.070      0.000
    SPACE              0.079      0.030      2.594      0.009
    HIGH_UTL           0.045      0.025      1.779      0.075

     Latent                                         Two-Tailed
    Variable        Estimate       S.E.  Est./S.E.    P-Value

    ERRBLF             0.175      0.047      3.759      0.000
    M_ADAP             0.239      0.058      4.119      0.000
    A_ADAP             0.211      0.051      4.097      0.000


STANDARDIZED TOTAL, TOTAL INDIRECT, SPECIFIC INDIRECT, AND DIRECT EFFECTS


STDYX Standardization

                                                    Two-Tailed
                    Estimate       S.E.  Est./S.E.    P-Value

Effects from GRBLF to SPACE

  Total                0.070      0.031      2.248      0.025
  Total indirect       0.070      0.031      2.248      0.025

  Specific indirect 1
    SPACE
    ERRBLF
    GRBLF             -0.002      0.004     -0.376      0.707

  Specific indirect 2
    SPACE
    M_ADAP
    GRBLF              0.008      0.030      0.266      0.790

  Specific indirect 3
    SPACE
    A_ADAP
    GRBLF              0.064      0.029      2.194      0.028


Effects from FXBLF to SPACE

  Total               -0.032      0.034     -0.951      0.342
  Total indirect      -0.032      0.034     -0.951      0.342

  Specific indirect 1
    SPACE
    ERRBLF
    FXBLF              0.016      0.019      0.843      0.399

  Specific indirect 2
    SPACE
    M_ADAP
    FXBLF             -0.002      0.010     -0.255      0.799

  Specific indirect 3
    SPACE
    A_ADAP
    FXBLF             -0.046      0.031     -1.494      0.135


Effects from GRBLF to HIGH_UTL

  Total                0.036      0.027      1.333      0.183
  Total indirect       0.036      0.027      1.333      0.183

  Specific indirect 1
    HIGH_UTL
    ERRBLF
    GRBLF              0.001      0.002      0.237      0.813

  Specific indirect 2
    HIGH_UTL
    M_ADAP
    GRBLF             -0.017      0.031     -0.552      0.581

  Specific indirect 3
    HIGH_UTL
    A_ADAP
    GRBLF              0.052      0.029      1.794      0.073


Effects from FXBLF to HIGH_UTL

  Total               -0.039      0.029     -1.357      0.175
  Total indirect      -0.039      0.029     -1.357      0.175

  Specific indirect 1
    HIGH_UTL
    ERRBLF
    FXBLF             -0.006      0.022     -0.275      0.783

  Specific indirect 2
    HIGH_UTL
    M_ADAP
    FXBLF              0.005      0.011      0.496      0.620

  Specific indirect 3
    HIGH_UTL
    A_ADAP
    FXBLF             -0.038      0.027     -1.403      0.161


CONFIDENCE INTERVALS OF STANDARDIZED MODEL RESULTS


STDYX Standardization

                  Lower .5%  Lower 2.5%    Lower 5%    Estimate    Upper 5%  Upper 2.5%   Upper .5%

 FXBLF    BY
    FXBLF1           0.320       0.358       0.377       0.478       0.579       0.598       0.636
    FXBLF2           0.430       0.464       0.482       0.572       0.663       0.680       0.714
    FXBLF3           0.350       0.387       0.406       0.504       0.603       0.621       0.658
    FXBLF4           0.345       0.383       0.403       0.505       0.608       0.627       0.666
    FXBLF5           0.529       0.564       0.583       0.678       0.773       0.791       0.827

 GRBLF    BY
    GRBLF1           0.644       0.668       0.680       0.743       0.806       0.818       0.842
    GRBLF2           0.747       0.765       0.773       0.819       0.865       0.874       0.891
    GRBLF3           0.683       0.705       0.716       0.774       0.833       0.844       0.866
    GRBLF4           0.828       0.840       0.847       0.880       0.914       0.920       0.933
    GRBLF5           0.795       0.811       0.819       0.860       0.902       0.910       0.925

 UNBLF    BY
    UNBLF1           0.738       0.757       0.768       0.821       0.874       0.884       0.904
    UNBLF2           0.759       0.776       0.785       0.831       0.878       0.887       0.904
    UNBLF3           0.784       0.801       0.809       0.854       0.898       0.907       0.923
    UNBLF4           0.712       0.729       0.739       0.786       0.833       0.843       0.860
    UNBLF5           0.798       0.813       0.820       0.860       0.900       0.907       0.922

 NUNBLF   BY
    NUNBLF1          0.518       0.545       0.558       0.630       0.702       0.715       0.742
    NUNBLF2          0.696       0.720       0.732       0.795       0.859       0.871       0.895
    NUNBLF3          0.622       0.654       0.670       0.756       0.842       0.859       0.891
    NUNBLF4          0.564       0.589       0.601       0.667       0.733       0.746       0.771
    NUNBLF5          0.600       0.623       0.635       0.697       0.759       0.771       0.795

 BRBLF    BY
    BRBLF1           0.646       0.666       0.677       0.732       0.786       0.797       0.817
    BRBLF2           0.552       0.575       0.586       0.647       0.708       0.720       0.742
    BRBLF3           0.810       0.823       0.829       0.864       0.899       0.905       0.918
    BRBLF4           0.796       0.810       0.817       0.855       0.893       0.900       0.914
    BRBLF5           0.812       0.824       0.830       0.863       0.895       0.902       0.914

 ERRBLF   BY
    ERRBLF1          0.484       0.510       0.524       0.594       0.664       0.677       0.704
    ERRBLF2          0.635       0.660       0.673       0.739       0.806       0.818       0.843
    ERRBLF3          0.733       0.755       0.766       0.824       0.883       0.894       0.916
    ERRBLF4          0.855       0.869       0.877       0.914       0.951       0.958       0.972
    ERRBLF5          0.779       0.797       0.806       0.853       0.901       0.910       0.928

 M_ADAP   BY
    M_ADAP1          0.445       0.475       0.491       0.572       0.653       0.668       0.699
    M_ADAP2          0.499       0.532       0.549       0.637       0.725       0.742       0.775
    M_ADAP3          0.431       0.466       0.484       0.577       0.670       0.687       0.722
    M_ADAP4          0.386       0.422       0.441       0.538       0.635       0.653       0.690
    M_ADAP5          0.469       0.508       0.527       0.630       0.734       0.753       0.792
    M_ADAP6          0.443       0.478       0.496       0.590       0.684       0.702       0.737

 A_ADAP   BY
    A_ADAP1          0.592       0.614       0.625       0.683       0.742       0.753       0.775
    A_ADAP2          0.399       0.427       0.441       0.516       0.590       0.605       0.633
    A_ADAP3          0.594       0.615       0.626       0.683       0.740       0.751       0.772
    A_ADAP4          0.587       0.614       0.628       0.701       0.774       0.788       0.815
    A_ADAP5          0.672       0.692       0.703       0.756       0.810       0.820       0.840
    A_ADAP6          0.482       0.510       0.524       0.599       0.674       0.688       0.716
    A_ADAP7          0.503       0.530       0.543       0.615       0.686       0.700       0.727

 ERRBLF   ON
    FXBLF           -0.550      -0.496      -0.469      -0.327      -0.186      -0.158      -0.105
    GRBLF           -0.171      -0.123      -0.098       0.032       0.161       0.186       0.235
    UNBLF           -0.141      -0.082      -0.052       0.106       0.264       0.294       0.353
    NUNBLF          -0.300      -0.204      -0.155       0.102       0.359       0.408       0.504
    BRBLF           -0.473      -0.396      -0.357      -0.151       0.055       0.094       0.171

 M_ADAP   ON
    FXBLF           -0.379      -0.313      -0.279      -0.104       0.072       0.105       0.171
    GRBLF            0.123       0.175       0.201       0.338       0.476       0.502       0.554
    UNBLF           -0.439      -0.378      -0.347      -0.183      -0.020       0.011       0.072
    NUNBLF          -0.552      -0.460      -0.413      -0.167       0.078       0.125       0.217
    BRBLF           -0.477      -0.398      -0.358      -0.148       0.062       0.102       0.181

 A_ADAP   ON
    FXBLF           -0.412      -0.352      -0.322      -0.164      -0.005       0.025       0.084
    GRBLF            0.033       0.080       0.103       0.226       0.349       0.373       0.419
    UNBLF           -0.093      -0.037      -0.009       0.140       0.288       0.317       0.372
    NUNBLF          -0.252      -0.156      -0.107       0.150       0.407       0.456       0.552
    BRBLF           -0.557      -0.478      -0.437      -0.226      -0.014       0.026       0.105

 SPACE    ON
    ERRBLF          -0.197      -0.162      -0.144      -0.050       0.044       0.062       0.098
    M_ADAP          -0.202      -0.148      -0.121       0.024       0.168       0.195       0.250
    A_ADAP           0.075       0.125       0.150       0.283       0.416       0.441       0.491

 HIGH_UTL ON
    ERRBLF          -0.152      -0.111      -0.090       0.019       0.127       0.148       0.189
    M_ADAP          -0.288      -0.231      -0.202      -0.051       0.101       0.130       0.186
    A_ADAP           0.001       0.056       0.084       0.231       0.379       0.407       0.462

 ERRBLF   WITH
    M_ADAP           0.054       0.098       0.120       0.238       0.356       0.378       0.423
    A_ADAP           0.134       0.175       0.196       0.305       0.415       0.436       0.477

 M_ADAP   WITH
    A_ADAP           0.380       0.423       0.444       0.557       0.670       0.691       0.733

 GRBLF    WITH
    FXBLF           -0.656      -0.618      -0.599      -0.497      -0.395      -0.375      -0.337

 UNBLF    WITH
    FXBLF           -0.461      -0.416      -0.393      -0.274      -0.155      -0.133      -0.088
    GRBLF            0.268       0.308       0.329       0.435       0.542       0.562       0.602

 NUNBLF   WITH
    FXBLF            0.214       0.259       0.283       0.405       0.527       0.550       0.596
    GRBLF           -0.459      -0.420      -0.400      -0.297      -0.194      -0.174      -0.135
    UNBLF           -0.760      -0.727      -0.710      -0.621      -0.532      -0.515      -0.481

 BRBLF    WITH
    FXBLF            0.247       0.284       0.304       0.405       0.505       0.525       0.562
    GRBLF           -0.371      -0.338      -0.321      -0.231      -0.141      -0.124      -0.091
    UNBLF           -0.605      -0.573      -0.556      -0.470      -0.383      -0.367      -0.334
    NUNBLF           0.716       0.737       0.748       0.803       0.858       0.868       0.889

 HIGH_UTL WITH
    SPACE            0.074       0.106       0.123       0.208       0.294       0.310       0.342

 Intercepts
    FXBLF1           1.726       1.762       1.781       1.877       1.973       1.992       2.028
    FXBLF2           1.962       2.003       2.024       2.134       2.244       2.265       2.307
    FXBLF3           1.702       1.738       1.756       1.850       1.945       1.963       1.998
    FXBLF4           1.676       1.706       1.721       1.801       1.881       1.896       1.926
    FXBLF5           1.956       2.002       2.025       2.148       2.271       2.295       2.341
    GRBLF1           4.294       4.478       4.571       5.062       5.552       5.646       5.829
    GRBLF2           4.093       4.278       4.373       4.867       5.362       5.456       5.641
    GRBLF3           4.931       5.199       5.336       6.052       6.767       6.904       7.172
    GRBLF4           4.400       4.572       4.660       5.121       5.581       5.669       5.842
    GRBLF5           4.065       4.237       4.326       4.786       5.247       5.335       5.507
    NUNBLF1          2.039       2.083       2.106       2.223       2.340       2.362       2.406
    NUNBLF2          1.687       1.719       1.735       1.820       1.906       1.922       1.954
    NUNBLF3          1.716       1.760       1.783       1.900       2.017       2.039       2.083
    NUNBLF4          1.985       2.030       2.052       2.171       2.290       2.313       2.357
    NUNBLF5          1.769       1.801       1.818       1.905       1.992       2.009       2.041
    UNBLF1           3.844       4.002       4.082       4.504       4.926       5.007       5.165
    UNBLF2           3.607       3.745       3.817       4.188       4.559       4.630       4.769
    UNBLF3           3.606       3.743       3.813       4.181       4.548       4.618       4.756
    UNBLF4           2.852       2.954       3.006       3.277       3.548       3.600       3.701
    UNBLF5           3.608       3.761       3.839       4.247       4.656       4.734       4.887
    BRBLF1           1.943       1.981       2.001       2.103       2.206       2.225       2.264
    BRBLF2           2.216       2.270       2.297       2.440       2.584       2.611       2.665
    BRBLF3           1.722       1.753       1.768       1.851       1.933       1.949       1.980
    BRBLF4           1.747       1.780       1.797       1.885       1.974       1.991       2.024
    BRBLF5           1.658       1.688       1.704       1.784       1.864       1.879       1.909
    ERRBLF1          6.795       7.099       7.255       8.070       8.884       9.040       9.345
    ERRBLF2          4.940       5.178       5.300       5.937       6.574       6.696       6.934
    ERRBLF3          4.777       4.992       5.102       5.676       6.251       6.361       6.576
    ERRBLF4          4.648       4.894       5.020       5.678       6.335       6.461       6.708
    ERRBLF5          4.899       5.168       5.306       6.024       6.742       6.879       7.148
    M_ADAP1          3.189       3.291       3.343       3.616       3.889       3.942       4.044
    M_ADAP2          2.749       2.822       2.859       3.054       3.250       3.287       3.360
    M_ADAP3          2.763       2.833       2.869       3.055       3.242       3.278       3.347
    M_ADAP4          2.261       2.309       2.334       2.462       2.590       2.615       2.662
    M_ADAP5          2.828       2.905       2.945       3.150       3.356       3.395       3.472
    M_ADAP6          3.424       3.551       3.616       3.955       4.293       4.358       4.485
    A_ADAP1          3.949       4.069       4.131       4.453       4.776       4.837       4.958
    A_ADAP2          4.897       5.089       5.187       5.701       6.215       6.313       6.506
    A_ADAP3          4.120       4.256       4.325       4.688       5.050       5.119       5.255
    A_ADAP4          5.098       5.318       5.431       6.019       6.607       6.720       6.940
    A_ADAP5          4.255       4.429       4.517       4.981       5.445       5.533       5.707
    A_ADAP6          4.717       4.940       5.055       5.652       6.250       6.364       6.588
    A_ADAP7          2.936       3.022       3.066       3.295       3.525       3.569       3.655
    SPACE            0.632       0.658       0.671       0.740       0.809       0.822       0.848
    HIGH_UTL         1.058       1.099       1.121       1.232       1.344       1.365       1.407

 Variances
    FXBLF            1.000       1.000       1.000       1.000       1.000       1.000       1.000
    GRBLF            1.000       1.000       1.000       1.000       1.000       1.000       1.000
    UNBLF            1.000       1.000       1.000       1.000       1.000       1.000       1.000
    NUNBLF           1.000       1.000       1.000       1.000       1.000       1.000       1.000
    BRBLF            1.000       1.000       1.000       1.000       1.000       1.000       1.000

 Residual Variances
    FXBLF1           0.621       0.657       0.675       0.772       0.868       0.886       0.922
    FXBLF2           0.511       0.549       0.569       0.673       0.776       0.796       0.835
    FXBLF3           0.590       0.628       0.647       0.746       0.845       0.864       0.901
    FXBLF4           0.582       0.621       0.641       0.745       0.848       0.868       0.907
    FXBLF5           0.339       0.387       0.412       0.541       0.670       0.694       0.743
    GRBLF1           0.301       0.336       0.354       0.448       0.542       0.560       0.595
    GRBLF2           0.210       0.239       0.253       0.329       0.404       0.418       0.447
    GRBLF3           0.259       0.292       0.310       0.400       0.491       0.508       0.542
    GRBLF4           0.132       0.154       0.166       0.225       0.284       0.295       0.318
    GRBLF5           0.148       0.175       0.189       0.260       0.331       0.345       0.372
    NUNBLF1          0.462       0.496       0.513       0.603       0.693       0.711       0.744
    NUNBLF2          0.210       0.247       0.267       0.368       0.469       0.488       0.526
    NUNBLF3          0.224       0.273       0.298       0.428       0.558       0.583       0.631
    NUNBLF4          0.417       0.450       0.467       0.555       0.643       0.660       0.693
    NUNBLF5          0.378       0.411       0.427       0.514       0.601       0.617       0.650
    UNBLF1           0.190       0.223       0.239       0.327       0.414       0.430       0.463
    UNBLF2           0.188       0.217       0.232       0.309       0.386       0.401       0.429
    UNBLF3           0.152       0.181       0.195       0.271       0.347       0.362       0.390
    UNBLF4           0.265       0.293       0.308       0.382       0.457       0.471       0.499
    UNBLF5           0.153       0.179       0.192       0.260       0.329       0.342       0.367
    BRBLF1           0.339       0.369       0.385       0.465       0.545       0.560       0.590
    BRBLF2           0.458       0.487       0.502       0.581       0.660       0.675       0.704
    BRBLF3           0.160       0.182       0.194       0.254       0.314       0.325       0.348
    BRBLF4           0.167       0.192       0.204       0.269       0.333       0.346       0.370
    BRBLF5           0.167       0.188       0.199       0.256       0.312       0.323       0.344
    ERRBLF1          0.517       0.548       0.564       0.647       0.731       0.747       0.778
    ERRBLF2          0.300       0.337       0.356       0.454       0.552       0.570       0.607
    ERRBLF3          0.170       0.206       0.224       0.320       0.417       0.435       0.471
    ERRBLF4          0.059       0.084       0.097       0.165       0.233       0.246       0.272
    ERRBLF5          0.144       0.174       0.190       0.272       0.353       0.369       0.399
    M_ADAP1          0.528       0.563       0.581       0.673       0.766       0.784       0.818
    M_ADAP2          0.419       0.461       0.482       0.594       0.706       0.728       0.770
    M_ADAP3          0.500       0.540       0.560       0.667       0.774       0.795       0.835
    M_ADAP4          0.547       0.586       0.606       0.711       0.815       0.835       0.874
    M_ADAP5          0.399       0.448       0.473       0.602       0.732       0.757       0.806
    M_ADAP6          0.478       0.520       0.541       0.652       0.763       0.784       0.826
    A_ADAP1          0.408       0.438       0.453       0.533       0.613       0.628       0.658
    A_ADAP2          0.613       0.642       0.657       0.734       0.811       0.826       0.855
    A_ADAP3          0.411       0.441       0.456       0.533       0.611       0.626       0.655
    A_ADAP4          0.348       0.386       0.406       0.508       0.610       0.630       0.668
    A_ADAP5          0.301       0.331       0.347       0.428       0.509       0.525       0.555
    A_ADAP6          0.501       0.535       0.552       0.641       0.731       0.748       0.781
    A_ADAP7          0.484       0.517       0.534       0.622       0.710       0.727       0.760
    SPACE            0.843       0.862       0.871       0.921       0.971       0.981       0.999
    HIGH_UTL         0.890       0.906       0.914       0.955       0.997       1.005       1.020
    ERRBLF           0.705       0.734       0.748       0.825       0.902       0.916       0.945
    M_ADAP           0.612       0.648       0.666       0.761       0.857       0.875       0.911
    A_ADAP           0.657       0.689       0.705       0.789       0.874       0.890       0.922

CONFIDENCE INTERVALS OF STANDARDIZED TOTAL, TOTAL INDIRECT, SPECIFIC INDIRECT, AND DIRECT EFFECTS


STDYX Standardization

                  Lower .5%  Lower 2.5%    Lower 5%    Estimate    Upper 5%  Upper 2.5%   Upper .5%

Effects from GRBLF to SPACE

  Total             -0.010       0.009       0.019       0.070       0.122       0.132       0.151
  Total indirect    -0.010       0.009       0.019       0.070       0.122       0.132       0.151

  Specific indirect 1
    SPACE
    ERRBLF
    GRBLF           -0.012      -0.010      -0.008      -0.002       0.005       0.007       0.009

  Specific indirect 2
    SPACE
    M_ADAP
    GRBLF           -0.069      -0.051      -0.041       0.008       0.057       0.067       0.085

  Specific indirect 3
    SPACE
    A_ADAP
    GRBLF           -0.011       0.007       0.016       0.064       0.112       0.121       0.139


Effects from FXBLF to SPACE

  Total             -0.120      -0.099      -0.089      -0.032       0.024       0.034       0.055
  Total indirect    -0.120      -0.099      -0.089      -0.032       0.024       0.034       0.055

  Specific indirect 1
    SPACE
    ERRBLF
    FXBLF           -0.034      -0.022      -0.016       0.016       0.048       0.054       0.066

  Specific indirect 2
    SPACE
    M_ADAP
    FXBLF           -0.027      -0.021      -0.018      -0.002       0.013       0.016       0.022

  Specific indirect 3
    SPACE
    A_ADAP
    FXBLF           -0.126      -0.107      -0.097      -0.046       0.005       0.014       0.034


Effects from GRBLF to HIGH_UTL

  Total             -0.033      -0.017      -0.008       0.036       0.080       0.088       0.105
  Total indirect    -0.033      -0.017      -0.008       0.036       0.080       0.088       0.105

  Specific indirect 1
    HIGH_UTL
    ERRBLF
    GRBLF           -0.006      -0.004      -0.004       0.001       0.005       0.005       0.007

  Specific indirect 2
    HIGH_UTL
    M_ADAP
    GRBLF           -0.097      -0.078      -0.068      -0.017       0.034       0.044       0.063

  Specific indirect 3
    HIGH_UTL
    A_ADAP
    GRBLF           -0.023      -0.005       0.004       0.052       0.100       0.110       0.128


Effects from FXBLF to HIGH_UTL

  Total             -0.112      -0.095      -0.086      -0.039       0.008       0.017       0.035
  Total indirect    -0.112      -0.095      -0.086      -0.039       0.008       0.017       0.035

  Specific indirect 1
    HIGH_UTL
    ERRBLF
    FXBLF           -0.063      -0.049      -0.042      -0.006       0.030       0.037       0.051

  Specific indirect 2
    HIGH_UTL
    M_ADAP
    FXBLF           -0.022      -0.016      -0.012       0.005       0.023       0.026       0.033

  Specific indirect 3
    HIGH_UTL
    A_ADAP
    FXBLF           -0.107      -0.091      -0.082      -0.038       0.007       0.015       0.032


MODEL MODIFICATION INDICES

NOTE:  Modification indices for direct effects of observed dependent variables
regressed on covariates may not be included.  To include these, request
MODINDICES (ALL).

Minimum M.I. value for printing the modification index    10.000

                                   M.I.     E.P.C.  Std E.P.C.  StdYX E.P.C.

BY Statements

GRBLF    BY A_ADAP1               13.687     0.274      0.175        0.194
GRBLF    BY A_ADAP4               10.887    -0.195     -0.125       -0.171
UNBLF    BY NUNBLF2               13.515     0.302      0.231        0.223
UNBLF    BY NUNBLF3               13.405     0.272      0.208        0.227
UNBLF    BY NUNBLF4               15.098    -0.438     -0.336       -0.256
UNBLF    BY A_ADAP4               12.335    -0.166     -0.127       -0.174
NUNBLF   BY BRBLF1                11.288     0.438      0.346        0.302
M_ADAP   BY A_ADAP1               28.507     0.563      0.358        0.398
A_ADAP   BY M_ADAP6               36.668     0.804      0.495        0.488

WITH Statements

FXBLF5   WITH FXBLF4              10.309     0.195      0.195        0.257
GRBLF2   WITH GRBLF1              28.068     0.109      0.109        0.374
GRBLF4   WITH GRBLF1              14.106    -0.069     -0.069       -0.301
GRBLF5   WITH GRBLF2              14.953    -0.076     -0.076       -0.328
GRBLF5   WITH GRBLF4              24.022     0.091      0.091        0.504
NUNBLF3  WITH NUNBLF2             82.267     0.278      0.278        0.736
NUNBLF4  WITH NUNBLF1             15.032     0.238      0.238        0.251
NUNBLF4  WITH NUNBLF3             21.544    -0.190     -0.190       -0.325
UNBLF1   WITH GRBLF5              10.944    -0.057     -0.057       -0.236
UNBLF3   WITH NUNBLF3             10.140     0.069      0.069        0.223
UNBLF4   WITH GRBLF3              10.660    -0.074     -0.074       -0.214
UNBLF4   WITH GRBLF5              15.179     0.091      0.091        0.273
UNBLF4   WITH UNBLF2              12.088    -0.101     -0.101       -0.253
BRBLF2   WITH BRBLF1              25.337     0.217      0.217        0.318
BRBLF5   WITH UNBLF1              12.261    -0.077     -0.077       -0.250
M_ADAP2  WITH M_ADAP1             23.412     0.280      0.280        0.342
M_ADAP3  WITH M_ADAP2             22.598     0.277      0.277        0.337
M_ADAP4  WITH M_ADAP2             18.364    -0.273     -0.273       -0.297
M_ADAP5  WITH M_ADAP4             40.729     0.383      0.383        0.439
A_ADAP1  WITH FXBLF4              10.474    -0.148     -0.148       -0.207
A_ADAP1  WITH M_ADAP6             28.211     0.185      0.185        0.344
A_ADAP4  WITH A_ADAP1             15.560    -0.095     -0.095       -0.278
A_ADAP5  WITH A_ADAP4             10.928     0.071      0.071        0.251
A_ADAP6  WITH A_ADAP3             10.112    -0.083     -0.083       -0.211
A_ADAP6  WITH A_ADAP4             24.542     0.106      0.106        0.333


TECHNICAL 4 OUTPUT


     ESTIMATES DERIVED FROM THE MODEL


           ESTIMATED MEANS FOR THE LATENT VARIABLES
              FXBLF         GRBLF         UNBLF         NUNBLF        BRBLF
              ________      ________      ________      ________      ________
                0.000         0.000         0.000         0.000         0.000


           ESTIMATED MEANS FOR THE LATENT VARIABLES
              ERRBLF        M_ADAP        A_ADAP        SPACE         HIGH_UTL
              ________      ________      ________      ________      ________
                0.000         0.000         0.000         0.353         0.603


           S.E. FOR ESTIMATED MEANS FOR THE LATENT VARIABLES
              FXBLF         GRBLF         UNBLF         NUNBLF        BRBLF
              ________      ________      ________      ________      ________
                0.000         0.000         0.000         0.000         0.000


           S.E. FOR ESTIMATED MEANS FOR THE LATENT VARIABLES
              ERRBLF        M_ADAP        A_ADAP        SPACE         HIGH_UTL
              ________      ________      ________      ________      ________
                0.000         0.000         0.000         0.026         0.026


           EST./S.E. FOR ESTIMATED MEANS FOR THE LATENT VARIABLES
              FXBLF         GRBLF         UNBLF         NUNBLF        BRBLF
              ________      ________      ________      ________      ________
                0.000         0.000         0.000         0.000         0.000


           EST./S.E. FOR ESTIMATED MEANS FOR THE LATENT VARIABLES
              ERRBLF        M_ADAP        A_ADAP        SPACE         HIGH_UTL
              ________      ________      ________      ________      ________
                0.000         0.000         0.000        13.592        22.887


           TWO-TAILED P-VALUE FOR ESTIMATED MEANS FOR THE LATENT VARIABLES
              FXBLF         GRBLF         UNBLF         NUNBLF        BRBLF
              ________      ________      ________      ________      ________
                1.000         1.000         1.000         1.000         1.000


           TWO-TAILED P-VALUE FOR ESTIMATED MEANS FOR THE LATENT VARIABLES
              ERRBLF        M_ADAP        A_ADAP        SPACE         HIGH_UTL
              ________      ________      ________      ________      ________
                1.000         1.000         1.000         0.000         0.000


           ESTIMATED COVARIANCE MATRIX FOR THE LATENT VARIABLES
              FXBLF         GRBLF         UNBLF         NUNBLF        BRBLF
              ________      ________      ________      ________      ________
 FXBLF          0.261
 GRBLF         -0.162         0.409
 UNBLF         -0.107         0.213         0.587
 NUNBLF         0.163        -0.150        -0.375         0.622
 BRBLF          0.173        -0.124        -0.302         0.531         0.704
 ERRBLF        -0.068         0.053         0.056        -0.061        -0.074
 M_ADAP        -0.114         0.160         0.081        -0.158        -0.169
 A_ADAP        -0.108         0.148         0.140        -0.122        -0.150
 SPACE         -0.021         0.032         0.028        -0.025        -0.031
 HIGH_UTL      -0.017         0.022         0.024        -0.018        -0.023


           ESTIMATED COVARIANCE MATRIX FOR THE LATENT VARIABLES
              ERRBLF        M_ADAP        A_ADAP        SPACE         HIGH_UTL
              ________      ________      ________      ________      ________
 ERRBLF         0.115
 M_ADAP         0.075         0.405
 A_ADAP         0.088         0.245         0.379
 SPACE          0.013         0.056         0.081         0.228
 HIGH_UTL       0.016         0.031         0.062         0.059         0.239


           S.E. FOR ESTIMATED COVARIANCE MATRIX FOR THE LATENT VARIABLES
              FXBLF         GRBLF         UNBLF         NUNBLF        BRBLF
              ________      ________      ________      ________      ________
 FXBLF          0.071
 GRBLF          0.031         0.066
 UNBLF          0.028         0.044         0.086
 NUNBLF         0.036         0.036         0.064         0.098
 BRBLF          0.037         0.032         0.050         0.066         0.084
 ERRBLF         0.018         0.014         0.015         0.017         0.019
 M_ADAP         0.031         0.031         0.036         0.043         0.044
 A_ADAP         0.028         0.028         0.032         0.039         0.034
 SPACE          0.009         0.009         0.009         0.010         0.010
 HIGH_UTL       0.008         0.009         0.008         0.010         0.010


           S.E. FOR ESTIMATED COVARIANCE MATRIX FOR THE LATENT VARIABLES
              ERRBLF        M_ADAP        A_ADAP        SPACE         HIGH_UTL
              ________      ________      ________      ________      ________
 ERRBLF         0.020
 M_ADAP         0.017         0.083
 A_ADAP         0.017         0.040         0.052
 SPACE          0.008         0.019         0.018         0.008
 HIGH_UTL       0.010         0.019         0.018         0.012         0.005


           EST./S.E. FOR ESTIMATED COVARIANCE MATRIX FOR THE LATENT VARIABLES
              FXBLF         GRBLF         UNBLF         NUNBLF        BRBLF
              ________      ________      ________      ________      ________
 FXBLF          3.657
 GRBLF         -5.187         6.159
 UNBLF         -3.808         4.866         6.844
 NUNBLF         4.565        -4.168        -5.893         6.341
 BRBLF          4.660        -3.845        -6.007         8.005         8.361
 ERRBLF        -3.877         3.808         3.806        -3.506        -3.943
 M_ADAP        -3.712         5.245         2.271        -3.649        -3.845
 A_ADAP        -3.860         5.341         4.388        -3.159        -4.368
 SPACE         -2.425         3.333         3.111        -2.436        -2.997
 HIGH_UTL      -2.148         2.425         2.857        -1.803        -2.218


           EST./S.E. FOR ESTIMATED COVARIANCE MATRIX FOR THE LATENT VARIABLES
              ERRBLF        M_ADAP        A_ADAP        SPACE         HIGH_UTL
              ________      ________      ________      ________      ________
 ERRBLF         5.811
 M_ADAP         4.371         4.913
 A_ADAP         5.207         6.159         7.347
 SPACE          1.502         2.940         4.645        29.704
 HIGH_UTL       1.706         1.613         3.536         4.955        44.161


           TWO-TAILED P-VALUE FOR ESTIMATED COVARIANCE MATRIX FOR THE LATENT VARIABLES
              FXBLF         GRBLF         UNBLF         NUNBLF        BRBLF
              ________      ________      ________      ________      ________
 FXBLF          0.000
 GRBLF          0.000         0.000
 UNBLF          0.000         0.000         0.000
 NUNBLF         0.000         0.000         0.000         0.000
 BRBLF          0.000         0.000         0.000         0.000         0.000
 ERRBLF         0.000         0.000         0.000         0.000         0.000
 M_ADAP         0.000         0.000         0.023         0.000         0.000
 A_ADAP         0.000         0.000         0.000         0.002         0.000
 SPACE          0.015         0.001         0.002         0.015         0.003
 HIGH_UTL       0.032         0.015         0.004         0.071         0.027


           TWO-TAILED P-VALUE FOR ESTIMATED COVARIANCE MATRIX FOR THE LATENT VARIABLES
              ERRBLF        M_ADAP        A_ADAP        SPACE         HIGH_UTL
              ________      ________      ________      ________      ________
 ERRBLF         0.000
 M_ADAP         0.000         0.000
 A_ADAP         0.000         0.000         0.000
 SPACE          0.133         0.003         0.000         0.000
 HIGH_UTL       0.088         0.107         0.000         0.000         0.000


           ESTIMATED CORRELATION MATRIX FOR THE LATENT VARIABLES
              FXBLF         GRBLF         UNBLF         NUNBLF        BRBLF
              ________      ________      ________      ________      ________
 FXBLF          1.000
 GRBLF         -0.497         1.000
 UNBLF         -0.274         0.435         1.000
 NUNBLF         0.405        -0.297        -0.621         1.000
 BRBLF          0.405        -0.231        -0.470         0.803         1.000
 ERRBLF        -0.392         0.245         0.217        -0.227        -0.259
 M_ADAP        -0.349         0.394         0.166        -0.315        -0.316
 A_ADAP        -0.345         0.376         0.296        -0.252        -0.290
 SPACE         -0.086         0.104         0.077        -0.067        -0.077
 HIGH_UTL      -0.069         0.072         0.064        -0.046        -0.056


           ESTIMATED CORRELATION MATRIX FOR THE LATENT VARIABLES
              ERRBLF        M_ADAP        A_ADAP        SPACE         HIGH_UTL
              ________      ________      ________      ________      ________
 ERRBLF         1.000
 M_ADAP         0.349         1.000
 A_ADAP         0.421         0.625         1.000
 SPACE          0.078         0.183         0.277         1.000
 HIGH_UTL       0.098         0.101         0.208         0.251         1.000


           S.E. FOR ESTIMATED CORRELATION MATRIX FOR THE LATENT VARIABLES
              FXBLF         GRBLF         UNBLF         NUNBLF        BRBLF
              ________      ________      ________      ________      ________
 FXBLF          0.000
 GRBLF          0.062         0.000
 UNBLF          0.072         0.065         0.000
 NUNBLF         0.074         0.063         0.054         0.000
 BRBLF          0.061         0.055         0.053         0.033         0.000
 ERRBLF         0.062         0.056         0.055         0.061         0.059
 M_ADAP         0.077         0.058         0.075         0.076         0.069
 A_ADAP         0.074         0.057         0.060         0.068         0.060
 SPACE          0.034         0.030         0.025         0.026         0.025
 HIGH_UTL       0.031         0.029         0.023         0.025         0.025


           S.E. FOR ESTIMATED CORRELATION MATRIX FOR THE LATENT VARIABLES
              ERRBLF        M_ADAP        A_ADAP        SPACE         HIGH_UTL
              ________      ________      ________      ________      ________
 ERRBLF         0.000
 M_ADAP         0.067         0.000
 A_ADAP         0.054         0.058         0.000
 SPACE          0.051         0.061         0.054         0.000
 HIGH_UTL       0.057         0.063         0.058         0.050         0.000


           EST./S.E. FOR ESTIMATED CORRELATION MATRIX FOR THE LATENT VARIABLES
              FXBLF         GRBLF         UNBLF         NUNBLF        BRBLF
              ________      ________      ________      ________      ________
 FXBLF        999.000
 GRBLF         -8.018       999.000
 UNBLF         -3.794         6.719       999.000
 NUNBLF         5.456        -4.730       -11.474       999.000
 BRBLF          6.601        -4.239        -8.927        24.001       999.000
 ERRBLF        -6.331         4.341         3.950        -3.728        -4.353
 M_ADAP        -4.523         6.763         2.204        -4.169        -4.565
 A_ADAP        -4.635         6.655         4.969        -3.674        -4.816
 SPACE         -2.527         3.395         3.114        -2.570        -3.050
 HIGH_UTL      -2.218         2.511         2.841        -1.873        -2.230


           EST./S.E. FOR ESTIMATED CORRELATION MATRIX FOR THE LATENT VARIABLES
              ERRBLF        M_ADAP        A_ADAP        SPACE         HIGH_UTL
              ________      ________      ________      ________      ________
 ERRBLF       999.000
 M_ADAP         5.221       999.000
 A_ADAP         7.844        10.803       999.000
 SPACE          1.529         3.001         5.088       999.000
 HIGH_UTL       1.724         1.591         3.596         5.037       999.000


           TWO-TAILED P-VALUE FOR ESTIMATED CORRELATION MATRIX FOR THE LATENT VARIABLES
              FXBLF         GRBLF         UNBLF         NUNBLF        BRBLF
              ________      ________      ________      ________      ________
 FXBLF          0.000
 GRBLF          0.000         0.000
 UNBLF          0.000         0.000         0.000
 NUNBLF         0.000         0.000         0.000         0.000
 BRBLF          0.000         0.000         0.000         0.000         0.000
 ERRBLF         0.000         0.000         0.000         0.000         0.000
 M_ADAP         0.000         0.000         0.028         0.000         0.000
 A_ADAP         0.000         0.000         0.000         0.000         0.000
 SPACE          0.011         0.001         0.002         0.010         0.002
 HIGH_UTL       0.027         0.012         0.004         0.061         0.026


           TWO-TAILED P-VALUE FOR ESTIMATED CORRELATION MATRIX FOR THE LATENT VARIABLES
              ERRBLF        M_ADAP        A_ADAP        SPACE         HIGH_UTL
              ________      ________      ________      ________      ________
 ERRBLF         0.000
 M_ADAP         0.000         0.000
 A_ADAP         0.000         0.000         0.000
 SPACE          0.126         0.003         0.000         0.000
 HIGH_UTL       0.085         0.112         0.000         0.000         0.000
```
